# Calcium dysregulation amplifies fibrotic responses to TGFβ in human Friedreich’s ataxia fibroblasts

**DOI:** 10.64898/2026.09.21.753166

**Authors:** Anna Stepanova, Hibiki Kawamata, Giovanni Manfredi

## Abstract

Friedreich’s ataxia (FA) is an inherited disease caused by loss of frataxin (*FXN*) and characterized by neurodegeneration and fatal cardiomyopathy. Cardiac fibrosis contributes to cardiomyopathy by stiffening the heart wall, yet the underlying mechanisms remain unknown. Here, we investigated pro-fibrotic predisposition in FA patient-derived fibroblasts, focusing on the role of cytosolic calcium (Ca) in TGFβ-driven fibroblast-to-myofibroblast transition (FMT). We found pro-fibrotic transcriptional priming in FA fibroblasts, alongside elevated expression of genes controlled by the Ca-responsive transcription factor NFAT. Upon FMT, FA myofibroblasts showed amplified induction of pro-fibrotic (*CCN2*, *NOX4*) and suppression of anti-fibrotic (*CCN3*) genes, which were inversely correlated with residual *FXN*. Mechanistically, FA fibroblasts exhibited elevated cytosolic Ca and strongly downregulated expression of the Na-Ca exchanger NCX1, which directly correlated with *FXN*. Furthermore, NCX1 inhibition in control fibroblasts recapitulated FA Ca phenotypes, whereas *NCX1* transduction in FA fibroblasts normalized Ca dynamics and blunted *CCN2* induction in FMT. These findings highlight NCX1 as a modulator of fibrotic reprogramming in FA and identify Ca dyshomeostasis as an intrinsic mechanism of fibrosis that could be targeted therapeutically.

## Introduction

Friedreich’s ataxia (FA) is an inherited autosomal-recessive multisystem disorder characterized by progressive neurodegeneration and lethal cardiomyopathy, which remains the leading cause of death (Tsou *et al*, 2011; Reetz *et al*, 2025). In most cases, FA stems from a GAA triplet expansion in the first intron of the frataxin (*FXN*) gene (Campuzano *et al*, 1996), causing a deficiency in the mitochondrial protein frataxin, which is involved in iron-sulfur cluster biosynthesis (Lill & Freibert, 2020). Disease severity is typically greater in patients with lower residual FXN levels, which depends on GAA expansion length (Li *et al*, 2015). In the FA heart, FXN deficiency leads to the accumulation of dysfunctional mitochondria in cardiomyocytes, myocardial necrosis, and extensive replacement fibrosis (Payne, 2022). As a consequence, collagen and other extracellular matrix (ECM) proteins accumulate within the heart’s interstitium, myocardial stiffening ensues, ultimately compromising cardiac function. Despite clinical knowledge of fibrotic signatures from cardiac MRI (Raman *et al*, 2011) and autoptic studies (Koeppen *et al*, 2020), the cellular and molecular mechanisms driving fibrosis in FA hearts remain largely unknown. This is a significant gap in knowledge because dysregulated pro-fibrotic process in tissues rich in fibroblasts, such as the heart, can contribute to disease pathogenesis.

The fibrotic process typically involves the transition of fibroblasts into activated myofibroblasts, a process critically regulated by transforming growth factor beta (TGFβ) and cytosolic calcium (Ca) levels (Baum & Duffy, 2011; Gibb *et al*, 2020). During the fibroblast-to-myofibroblast transition (FMT), cells gain a contractile phenotype, express α-smooth muscle actin (αSMA), and increase the production of ECM components (Tai *et al*, 2021). Pro-fibrotic factors, such as TGFβ and mechanical stress, contribute to fibroblast activation and transcriptional control of fibrotic genes through SMAD-dependent and SMAD-independent pathways (Frangogiannis, 2020). Research increasingly suggests that fibroblast signaling maintains and escalates the fibrotic process through autocrine and paracrine loops (Miyara *et al*, 2025). In FMT, Ca acts as an intermediary in TGFβ and mechano-sensing signaling, and previous studies have shown that sustained elevation of cytosolic Ca levels augments the production of the pro-fibrotic cellular communication network factor 2 (*CCN2*) (Romero *et al*, 2005). A rise in cytosolic Ca can be triggered through various plasma membrane channels, such as Transient Receptor Potential Canonical 3 (TRPC3), Piezo1 (Li *et al*, 2026), TRPV4 (Adapala *et al*, 2013) or Orai1 (Zhang *et al*, 2016). It activates fibrotic transcriptional programs through multiple pathways, including Ca/calmodulin-dependent protein kinase II (CaMKII) (Janssen *et al*, 2015) and Nuclear Factor of Activated T cells (NFAT) (Saliba *et al*, 2019). This evidence indicates that cytosolic Ca dysregulation is a pro-fibrotic stimulus that could be targeted for therapeutic modulation. However, whether fibrosis associated with FA cardiomyopathy is linked to Ca dyshomeostasis remains to be determined. While studies in FXN-deficient mouse cardiomyocytes, cerebellar granule neurons (Abeti *et al*, 2018b), and astrocytes (Marullo *et al*, 2025) show altered Ca homeostasis, this has not been studied in FA human fibroblasts.

Understanding how fibroblast activation and Ca-related signaling pathways contribute to fibrosis may help identify therapeutic targets to control fibrotic progression in FA. We therefore asked whether human FA fibroblasts display intrinsic pro-fibrotic properties and alterations in cytosolic Ca handling in relation to FMT. To answer this question, we employed a cohort of human primary skin fibroblasts with graded GAA repeat lengths. This *in vitro* model of FA recapitulates the genetic lesion and the effects of varying FXN levels. We integrated bulk transcriptomic and proteomic profiling with functional assessment of cytosolic Ca handling to characterize the molecular and physiological phenotype of human FA fibroblasts during TGFβ-induced FMT.

## Results

### FA fibroblasts exhibit profibrotic gene expression profiles and sensitization to TGFβ signaling

To comprehensively investigate the molecular phenotype of FA patient fibroblasts, we performed bulk RNA sequencing on 12 FA lines and 12 healthy controls. For each group, we analyzed 6 females and 6 males. This fibroblast cohort is a subset of a previously reported FA fibroblast cohort (Li *et al*, 2016). FA patient lines investigated here had GAA short-allele repeats ranging from 367-916 and long-allele repeats ranging from 556-1470, and age at the time of skin biopsy ranged between 10 and 40 years in both groups. Differential expression gene (DEG) analysis identified 79 upregulated and 58 downregulated genes in FA relative to controls (p-adj < 0.05, Fig. 1A). As expected, *FXN* was significantly downregulated in patient cells. Notably, we observed reduced expression of *CCN2* (encoding a profibrotic cytokine) and *SLC8A1* (encoding the sodium-Ca exchanger NCX1).

**Figure 1.**
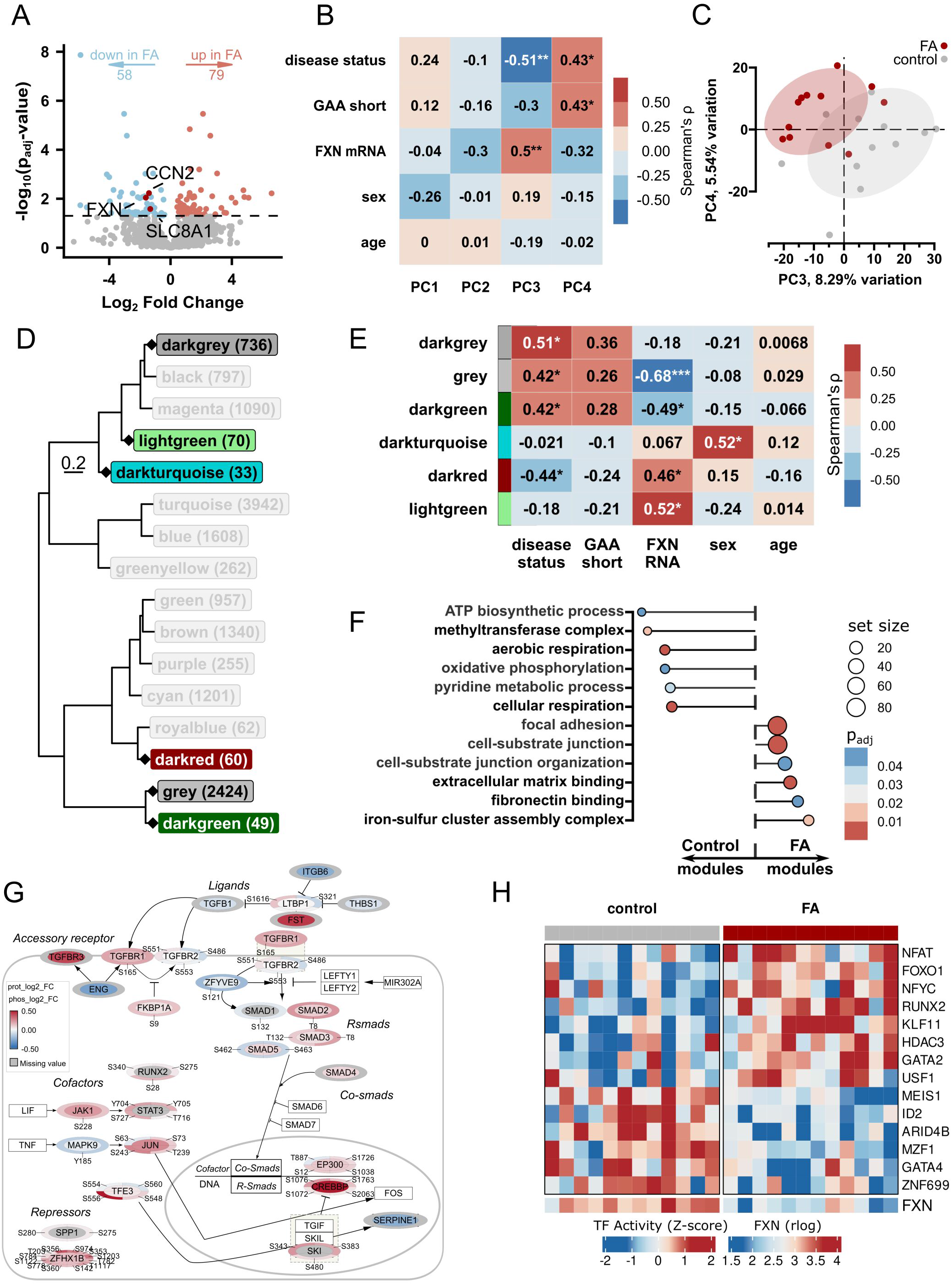
FA fibroblasts exhibit profibrotic gene expression profiles and sensitization to TGFβ signaling. A. Volcano plot of differential gene expression comparing patient (FA) versus control fibroblasts (n=12 cell lines per group). The negative log_10_ of the p_adj_-value (-log_10_(p_adj_-value)) is plotted against the log_2_ fold-change (log_2_FC). Genes passing the significance threshold (p_adj_-value < 0.05, dashed line) are colored by upregulation (red) or downregulation (blue) in FA cells. Key genes *FXN*, *CCN2*, and *SLC8A1* (NCX1) are highlighted and labeled. The total number of significantly down- and up-regulated genes is indicated at the top. B. Heatmap showing Spearman’s rank correlation coefficient (ρ) between the first four principal components (PC1-PC4) of the bulk transcriptome and sample metadata: disease status (FA or control), GAA short repeat length, *FXN* mRNA expression, sex, and age. Each cell shows the correlation coefficient; n = 24 total lines. C. Principal component analysis (PCA) bi-plot of sample clustering of PC3 versus PC4, with samples colored by group (control in grey, FA in red, n=12 cell lines per group). Ellipses encompass 75% of the data points for each group, illustrating the separation of FA and control samples along these components. D. Hierarchical clustering dendrogram of module eigengenes (ME) from WGCNA module dendrogram (n = 23 total samples, as one sample did not pass the quality threshold for the analysis). Each leaf represents a co-expression module, and the branch length (scale bar of 0.2 is shown) indicates the dissimilarity between modules. Modules of interest that correlated with patient traits (darkturquoise, darkred, darkgreen, darkgrey, grey, and lightgreen) are highlighted with their names and the number of genes in parentheses. E. WGCNA module-trait correlation heatmap showing Spearman’s rank correlation (ρ) and associated p-value (asterisks) between the ME (rows) and the sample traits (columns, n = 23 total samples). Color intensity represents the correlation strength, with red indicating a positive correlation and blue indicating a negative one. F. Gene Ontology (GO) enrichment comparison of WGCNA modules. Dot plot showing the top enriched GO Biological Process terms for modules highly correlated with controls (negative Fold Enrichment, darkturquoise and lightgreen modules) or FA (positive Fold Enrichment, darkred and darkgreen modules). Circle size represents the number of genes in the term, and color intensity represents the p_adj_-value. G. Network visualization of proteomics and phosphoproteomics mapped onto the simplified WikiPathways (WP:WP560) “TGF-beta Receptor Signaling” (FA versus control, n=8 cell lines per group). Node color (ellipses) represents the log_2_ fold-change of the protein level. Donut chart section color visualizes the log_2_ fold-change of each phosphorylation site abundance. T indicates phosphorylated threonine and S serine. Missing values are shown in grey. Proteins not identified by either proteomics or phosphoproteomics are shown in clear rectangles. H. Heatmap of predicted transcription factor (TF) activities significantly correlated (p < 0.05) with FXN mRNA expression (rlog counts) in control and FA fibroblasts (n = 12 cell lines per group). Data information: In B and E, asterisks indicate significance of Spearman’s rank correlation: * p < 0.05, ** p < 0.01, *** p < 0.001. In A and F, p_adj_-value are calculated using the Benjamini-Hochberg false discovery rate (FDR) with a significance threshold of p_adj_-value < 0.05.

Given the modest number of differentially expressed genes, we applied dimensionality reduction and network approaches to examine broad transcriptional patterns. Principal component (PC) analysis revealed that disease-associated variance was captured primarily by PC3 and PC4, which together explained 12-14% of the total variance (Fig. 1B). These components correlated significantly with disease status (FA or control), short-allele GAA repeat length, and *FXN* mRNA levels. When samples were plotted along these axes (PC3 vs. PC4), FA and control fibroblasts formed distinct clusters, with FA samples showing notably tighter clustering, suggesting lower transcriptional variability among FA lines than controls (Fig. 1C).

To identify coordinated gene expression programs, we performed weighted gene co-expression network analysis (WGCNA), which clusters genes into modules based on correlated expression patterns. Sixteen gene modules were identified (Fig. 1D). Of these, three modules (darkgray, gray, and darkgreen) were positively correlated with FA status and negatively correlated with *FXN* expression (Fig. 1E). Gene Ontology enrichment revealed that these three FA modules were enriched for extracellular matrix binding, focal adhesion, cell-substrate junctions, and fibronectin binding terms (Fig. 1F), which are associated with fibrotic programs. On the other hand, two control modules (darkred and lightgreen) were enriched for biosynthetic processes, oxidative phosphorylation, and cellular respiration.

To extend these FA-related transcriptional findings to the protein level, we performed TMT-based proteomics and TiO_2_-enriched phosphoproteomics on a randomly selected subset of 8 control and 8 FA fibroblast lines. To investigate canonical TGFβ-mediated fibrotic processes, we mapped the proteomics and phosphoproteomics data onto the WikiPathways “TGF-beta Receptor Signaling” (WP: WP560). We observed that endogenous TGFβ1 was downregulated in FA fibroblasts, yet downstream signaling components, including TGFβ receptors (TGFBR1 and TGFBR3) and SMAD2/3, showed increased protein abundance (Fig. 1G). Furthermore, SMAD2/3 showed increased activating phosphorylation marks, such as phosphorylation of threonine 8 (SMAD2/3 T8, Fig. 1G). Together, these data suggest that in FA fibroblasts, the TGFβ intracellular signaling components are enhanced, but low basal TGFβ1 ligand levels limit the activation of the pathway.

Next, to understand the upstream transcriptional regulatory mechanisms underlying profibrotic priming in FA fibroblasts, we used bioinformatic approaches to predict transcription factor (TF) activities from the gene expression data. For all samples, we evaluated TF activity scores and then compared FA and control groups. Statistically significant TFs were correlated with *FXN* gene expression, and only significantly correlated TFs were considered. Among the TFs predicted to be more active in FA, we identified NFAT (Fig. 1H), a Ca-calmodulin-responsive TF implicated in TGFβ-mediated FMT (Gibb *et al*, 2020). Another notable TF that was predicted to be more active in FA was RUNX2, which plays a profibrotic role as a cofactor of SMADs (Wu *et al*, 2026) and could act synergistically with NFAT. Interestingly, RUNX2 activity was linked to intracellular Ca levels in mesenchymal cells (Fromigué *et al*, 2010).

Together, these multi-omic data suggest that *FXN* deficiency alters intrinsic pro-fibrotic programs in FA fibroblasts at baseline low levels of endogenous intracellular TGFβ.

### TGFβ-induced FMT drives fibrotic reprogramming

The differences in the expression of key TGFβ signaling players observed in FA fibroblasts prompted us to test the effects of exogenous TGFβ stimulation. We subjected control and FA fibroblasts to a well-established fibroblast-to-myofibroblast differentiation protocol, which involves overnight serum starvation (0.5% FBS) followed by treatment with TGFβ1 (10 ng/mL in 1% FBS media) for 4-6 days (Piersma *et al*, 2017) (Fig. 2A).

**Figure 2.**
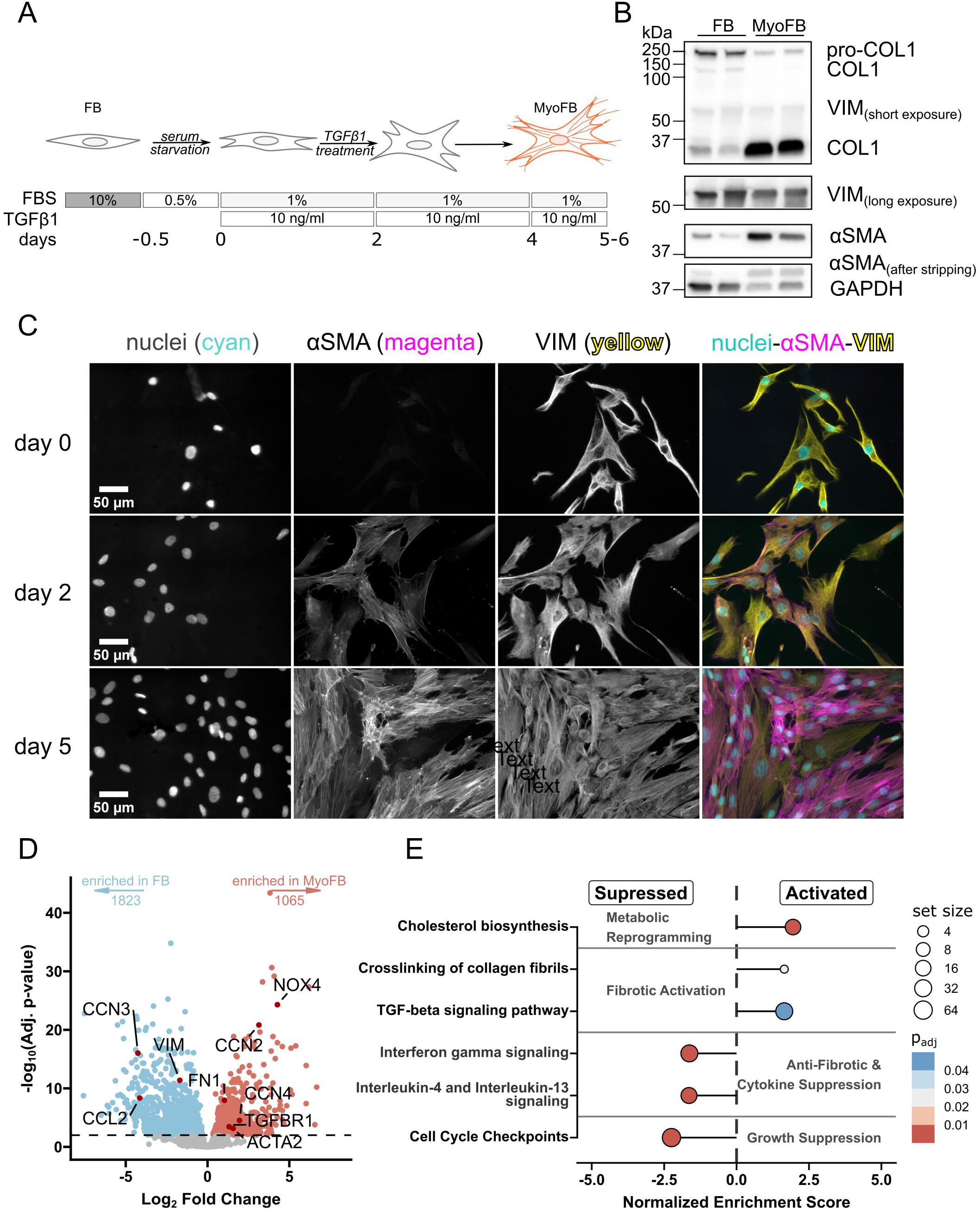
TGFβ-induced fibroblast-to-myofibroblast transition (FTM) drives fibrotic reprogramming. A. Schematic representation of the fibroblast-to-myofibroblast transition (FMT) protocol, indicating media changes and TGFβ1 treatment. Human skin fibroblasts (FB) were cultured under serum starvation conditions (0.5% FBS overnight) and stimulated with TGFβ1 (10 ng/ml) for at least 72 hours in reduced serum media (1% FBS) to induce differentiation into myofibroblasts (MyoFB). B. Western blot of myofibroblast differentiation markers in FB and MyoFB: ɑSMA (ɑ-smooth muscle actin), pro-Collagen (∼ 150 kDa), and COL1 bands (collagen I, ∼130 kDa and ∼ 35 kDa). VIM (vimentin), a general mesenchymal marker, and GAPDH, as the loading control (n = 2 lines per group). C. Immunocytochemistry staining of FB (Day 0) and MyoFB (Day 5). Cells were stained for ɑSMA (orange), vimentin (green), and the nuclear counterstain DAPI (blue) to illustrate cytoskeletal reorganization and ɑSMA-positive stress fiber formation. Scale bar: 50 μm. D. Volcano plot of transcriptomic changes during FMT comparing MyoFB versus FB (controls and FA combined, n = 24 cell lines per group). The vertical axis is the negative log_10_ of the p_adj_-value, and the horizontal axis is the log_2_FC. Enriched genes are colored in blue (enriched in FB) or red (enriched in MyoFB). Genes associated with myofibroblast differentiation (*ACTA2*, *COL1A1*, *FN1*, and *TGFBR1*, *CCN2*, *CCN4*, *NOX4* and *CCL2*) are highlighted and labeled. The total number of significantly enriched genes (Adj. p-value <0.01) is indicated at the top. E. Gene set enrichment analysis (GSEA) of FMT showing curated pathways from KEGG and Reactome grouped into metabolic reprogramming, fibrotic activation, anti-fibrotic and cytokine suppression, and growth suppression. The horizontal axis shows the normalized enrichment score (NES), where a positive NES indicates pathway activation (enriched in MyoFB) and a negative NES indicates pathway suppression (enriched in FB). Dot size is the number of genes from the core enrichment list within the pathway. Dot color represents the p_adj_-value.

This protocol models the FMT that occurs during tissue injury and fibrosis, a process highly relevant to cardiac pathology in FA.

We confirmed successful myofibroblast differentiation at multiple levels. Western blot analysis demonstrated robust upregulation of the canonical myofibroblast protein markers, α-smooth muscle actin (αSMA), pro-collagen I, and mature collagen I (Fig. 2B, Appendix Fig. S1). Immunofluorescence staining revealed the expected cytoskeletal reorganization in virtually all cells by day 5, with prominent αSMA-positive stress fibers and a decreased vimentin network, characteristic of the contractile myofibroblast phenotype (Fig. 2C). These molecular changes were similar in control and FA cells, indicating that FXN-deficient cells differentiate into myofibroblasts like controls.

To profile the transcriptional programs underlying FMT, we performed RNA sequencing on all 24 cell lines (12 FA and 12 controls), before and after TGFβ treatment, and compared gene expression in all fibroblasts and myofibroblasts, regardless of genotype. In cells treated with TGFβ, we observed extensive transcriptional remodeling associated with differentiation into myofibroblasts, with 1065 upregulated and 1823 downregulated genes (p-adj < 0.01, Fig. 2D). Upregulated genes included the expected myofibroblast markers, such as *ACTA2* (encoding αSMA), *COL1A1*, and *FN1*, as well as profibrotic mediators including *TGFBR1*, *CCN2*, *CCN4*, and *NOX4*. In addition, anti-fibrotic cytokines *CCL2* and *CCN3* were downregulated. Gene set enrichment analysis also revealed the coordinated biological programs involved in FMT. In myofibroblasts, we observed activation of cholesterol biosynthesis, TGFβ signaling cascades, and collagen fibril crosslinking pathways (Fig. 2E). In contrast, pathways associated with cell proliferation and growth were suppressed, as were anti-fibrotic cytokine programs. This data indicates that a transcriptional shift from a proliferative, growth-oriented state to a biosynthetic, matrix-producing state occurred. The coordinated suppression of growth pathways, alongside the activation of ECM production programs, defines the myofibroblast phenotype and sets the stage for examining disease-specific differences in this activated state.

### FA fibroblasts display an exacerbated fibrotic response upon TGFβ1-induced transition to myofibroblasts

Having established that both control and FA cells are equally capable of transforming into myofibroblasts upon stimulation with TGFβ1, we next asked whether FA myofibroblasts exhibit genotype-specific alterations. RNA sequencing comparison of 12 control and 12 FA myofibroblast lines revealed 29 upregulated and 56 downregulated genes (Fig. 3A), of which only a minority, 9 and 11, respectively, were in common between myofibroblasts and fibroblasts (Fig. EV1A). As expected, *FXN* expression levels were significantly downregulated, confirming persistent FXN deficiency. Among genes upregulated specifically in FA myofibroblasts, *GJA1* (encoding Connexin 43) was particularly notable, since this gap junction protein mediates intercellular communication and, in cardiac tissue, enables fibroblast-cardiomyocyte coupling that can affect electrical conductance (McArthur *et al*, 2015). Connexin 43 alterations have also been reported in post-mortem human FA heart (Koeppen *et al*, 2016a) and dorsal root ganglia (Koeppen *et al*, 2016b). Among the DEGs in common between fibroblasts and myofibroblasts, we identified a few transcripts associated with fibrotic programs: *KRT8*, *KRT18*, and *HAND2*-*AS1* (Fig. EV1A). Interestingly, *HAND2-AS1* is involved in fine-tuning epithelial-to-mesenchymal transition, and its downregulation is associated with an increased TGFβ1-induced profibrotic response (Vazana-Netzaíim *eī al*, 2023).

**Figure 3.**
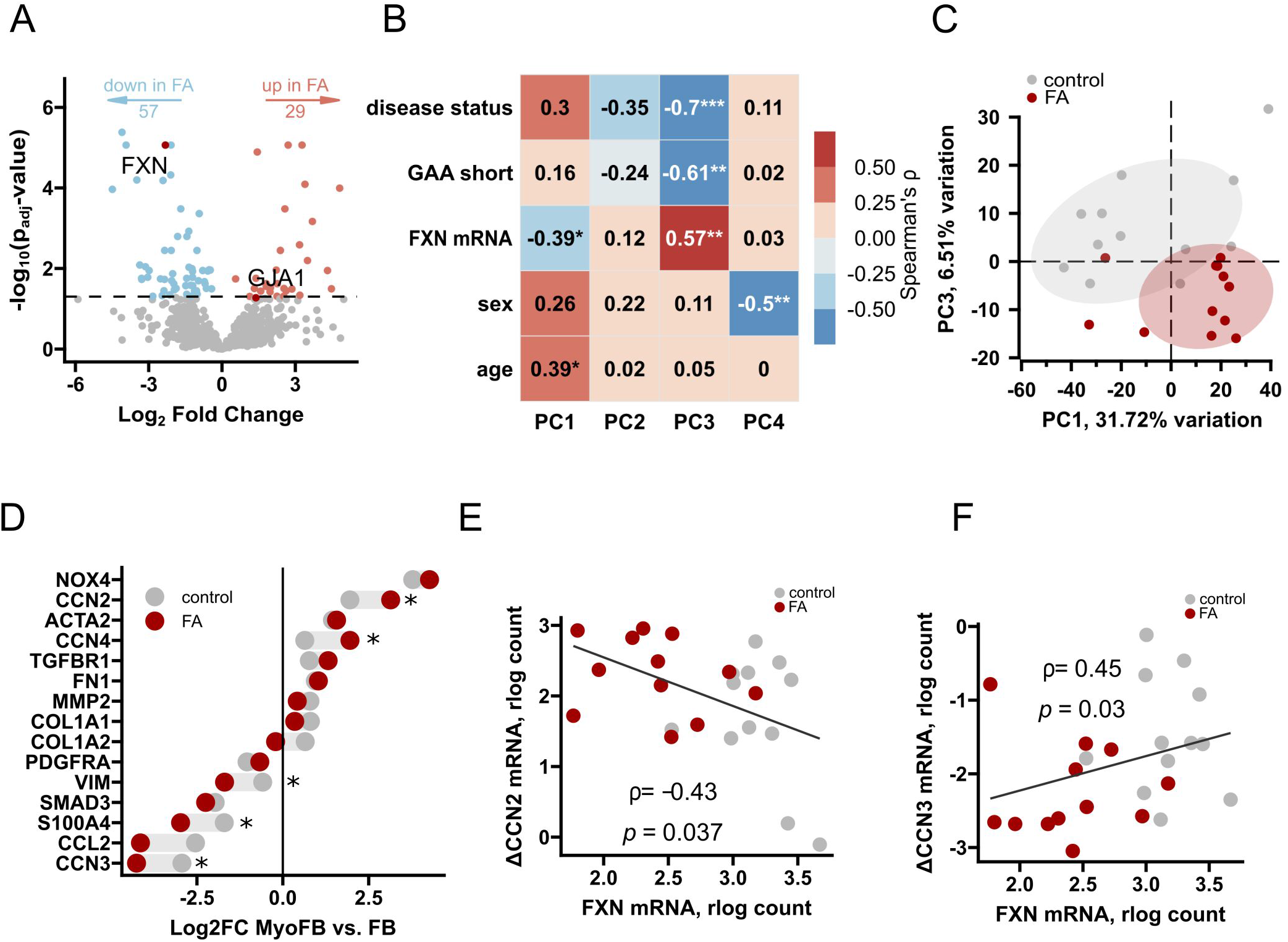
FA fibroblasts display an exacerbated fibrotic response to TGFβ1, which is inversely correlated with frataxin levels. A. Volcano plot of differential gene expression comparing patient (FA) versus control myofibroblasts (n=12 cell lines per group). Negative log_10_ of the p_adj_-value (-log_10_(p_adj_-value)) is plotted against the log_2_FC. Genes passing the significance threshold (p_adj_-value < 0.05, dashed line) are colored by upregulation (red) or downregulation (blue) in FA cells. Key genes *FXN* and *GJA1* are highlighted and labeled. The total number of significantly down- and up-regulated genes is indicated at the top. B. Heatmap showing Spearman’s rank correlation coefficient (ρ) between the first four principal components (PC1-PC4) of the bulk transcriptome and sample metadata: disease status (FA or control), GAA short repeat length, FXN mRNA expression, sex, and age in MyoFB. Each cell shows the correlation coefficient; n = 24 total cell lines. C. PCA bi-plot of sample clustering of PC1 versus PC3, with samples colored by group (control in grey, FA in red, n=12 cell lines per group). Ellipses encompass 75% of the data points for each group, illustrating the separation of FA and control samples along these components. D. Connected dot plot comparing the log_2_FC of MyoFB versus FB expression for a curated list of fibrotic genes, segregated by control (grey) and FA (red) genotypes. Asterisks mark changes that were significantly (p<0.05) different between control and FA upon FMT. E, F. Scatter plots showing the significant moderate Spearman’s rank correlation between the baseline *FXN* mRNA level (rlog count in FB) and the change in *CCN2* (E) or *CCN3* (F) mRNA expression (ΔCCN2/3 = CCN2/3_MyoFB_ - CCN2/3_FB_) upon FMT (n = 24 total samples). Data information: In B, E, and F, asterisks indicate the significance of Spearman’s rank correlation: * p < 0.05, ** p < 0.01, *** p < 0.001. In A and D, p_adj_-value are calculated using the Benjamini-Hochberg false discovery rate (FDR) with a significance threshold of p_adj_-value < 0.05. In D, the interaction term of a two-way ANOVA DESeq2 design (*∼ cell type + genotype + genotype:cell type*) was used to identify significant genes.

Principal component analysis showed that PC3, which explains 6.5% of variance, was negatively correlated with disease status and positively correlated with *FXN* expression (Fig. 3B). Plotting samples along PC1 and PC3 revealed separation between control and FA myofibroblasts, with FA samples forming a tighter cluster than controls (Fig. 3C), a pattern consistent with our observations in FA fibroblasts (Fig. 1C).

To specifically examine fibrotic programming, we used systematic literature mining and STRING database analysis to curate a consensus panel of fibrosis-associated genes with established roles in TGFβ signaling, ECM remodeling, and myofibroblast biology. To analyze genotype-specific differences in how cells respond to TGFβ differentiation signals, we calculated the fold change in expression during FMT (ΔGene = Gene_MyoFB_ -Gene_FB_) for each cell line individually. Analysis of “fibrotic” genes revealed striking genotype-dependent patterns. Profibrotic genes, including *NOX4*, *CCN2*, and *CCN4*, showed greater induction in FA cells than in controls. The most pronounced increase in induction was observed for *CCN2*, an early TGFβ-responsive gene and master regulator of fibrosis. Conversely, *CCN3*, considered an anti-fibrotic factor, exhibited greater suppression in FA myofibroblasts (Fig. 3D).

To determine whether there was a relationship between the magnitude of fibrotic gene expression response and *FXN*, we examined the correlation between ΔGene and *FXN* mRNA levels in fibroblasts and found that *ΔCCN2* was negatively correlated with *FXN*, indicating that lower *FXN* expression predicted greater *CCN2* induction upon TGFβ stimulation (Fig. 3E). In contrast, *ΔCCN3* showed a positive correlation with *FXN*, indicating that lower *FXN* expression predicted greater *CCN3* suppression upon TGFβ stimulation (Fig. 3F). These correlations suggest that *FXN* levels define profibrotic and anti-fibrotic gene expression changes in response to TGFβ stimulation in fibroblasts, and that FA fibroblasts mount amplified fibrotic responses to TGFβ stimulation, which inversely correlate with residual *FXN* levels.

### FA fibroblasts show altered cytosolic Ca homeostasis

It has been proposed that cytosolic Ca levels in fibroblasts influence their transformation to myofibroblasts (Gibb *et al*, 2020). Specifically, elevated cytosolic Ca levels promote profibrotic signaling. However, whether Ca homeostasis is altered in FA fibroblasts is unknown. Cytosolic Ca is tightly regulated, and basal Ca concentration is typically low, in the sub-micromolar range. When Ca is released from the endoplasmic reticulum (ER), the major intracellular Ca store, several mechanisms rapidly clear Ca from the cytosol, mainly plasma membrane extrusion, ER reuptake, and mitochondrial uptake.

To study cytosolic Ca homeostasis in FA fibroblasts, we used a genetically encoded Ca indicator, GCaMP6f, localized to the cytosol and measured Ca, both at baseline and upon ER Ca release induced by histamine (Fig. 4A, B). At the end of the experiment, cells were permeabilized with digitonin and treated with a saturating Ca concentration to establish the maximal fluorescence for data normalization. Comparison of cell populations from 12 control and 12 FA fibroblast lines revealed small but significant increases in baseline Ca (Fig. 4B, C), histamine-induced Ca peak amplitude (Fig. 4B, D), as well as the area under the curve (AUC, Fig. 4B, E) in FA fibroblasts. These findings suggest altered regulation of cytosolic Ca in FA fibroblasts.

**Figure 4.**
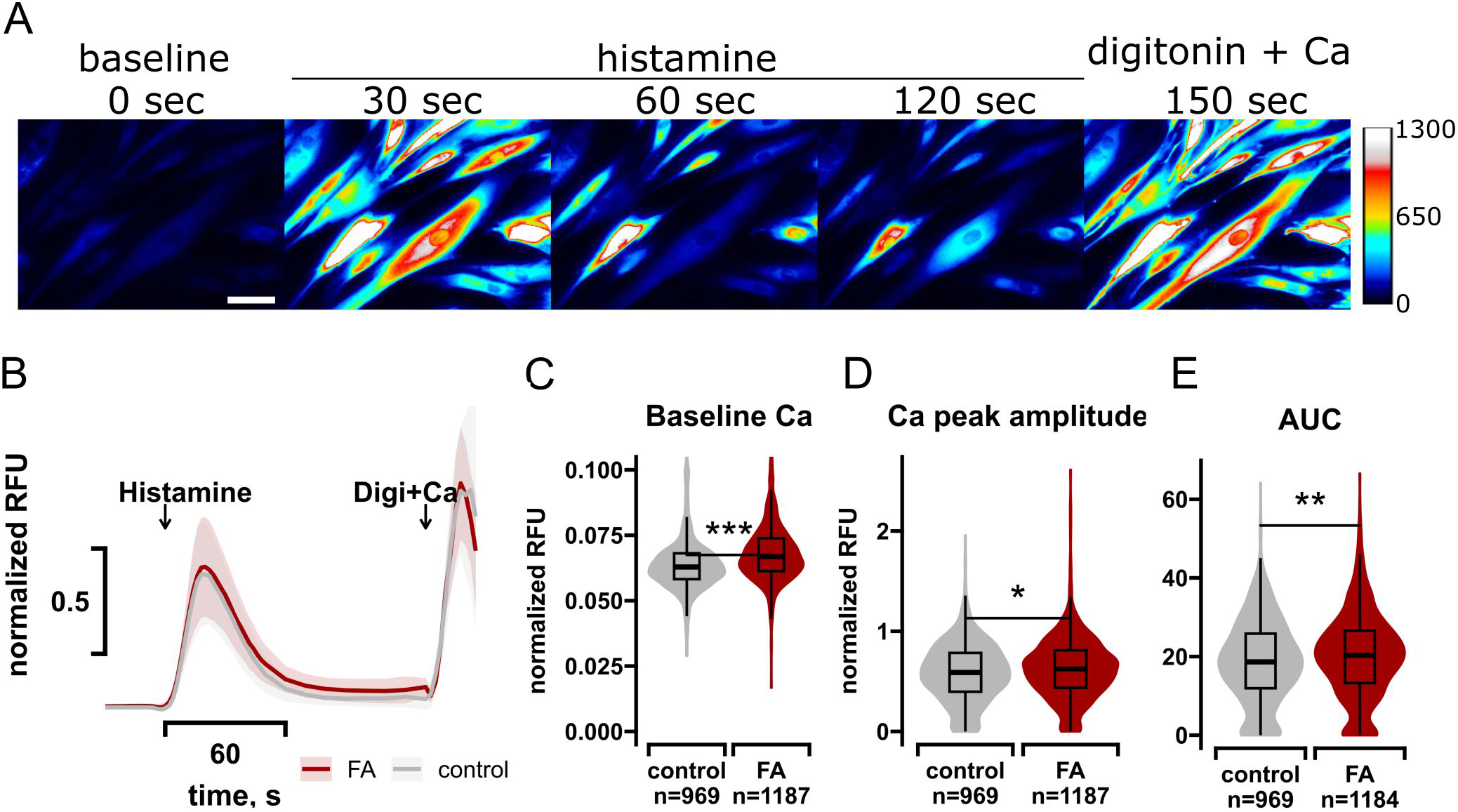
FA fibroblasts show altered cytosolic Ca homeostasis. A. Representative pseudocolor live-cell GCaMP6f images of cytosolic Ca in control and FA fibroblasts at baseline, during histamine stimulation (100 μM), and after addition of digitonin (10 μM) and Ca (10 mM). Fluorescence intensity is encoded using a “royal” color map, with warmer colors indicating higher GCaMP6f signal and thus higher cytosolic Ca levels. The times at which each representative image was taken are indicated. Scale bar: 50 µm. B. Traces of normalized GCaMP fluorescence ratio (RFU) over time. Arrows indicate addition of histamine, followed by digitonin (Digi) plus Ca, for normalization. The solid line shows the median trace for each genotype (n = ∼ 1000 cells per group from 12 cell lines per group), and the shaded areas indicate variability represented by the median absolute deviation (MAD). C-E. Quantification of histamine-induced Ca parameters in individual cells. Violin plots show the distribution and density of baseline cytosolic Ca (C), histamine-evoked peak amplitude (D), and area under the curve (AUC) for the primary histamine-induced peak (E), with internal boxplots indicating the median and interquartile range. The number of cells measured is indicated for each group (from 12 cell lines per group). Data information: In C-E, asterisks indicate significance of Wilcoxon test: * p < 0.05, ** p < 0.01, *** p < 0.001.

### Mitochondrial bioenergetics does not contribute to cytosolic Ca alterations in FA fibroblasts

Altered cytosolic Ca homeostasis can be the result of impaired mitochondrial membrane potential, which drives mitochondrial Ca uptake. FXN plays essential roles in FeS cluster assembly, and its deficiency may cause mitochondrial dysfunction, including decreased activities of FeS cluster-containing enzymes, such as respiratory chain complexes I and II and aconitase. This could impair mitochondrial respiration and membrane potential. Therefore, we assessed mitochondrial function to determine if respiratory and membrane potential impairments are associated with Ca dyshomeostasis in FA fibroblasts and myofibroblasts.

First, we quantified the enzymatic activities of complexes I and II in a randomly selected subset of control and FA fibroblasts and myofibroblasts (n=4-6). Complex I activity was measured as rotenone-sensitive NADH:decylubiquinone oxidoreductase activity, and complex II activity was assessed as atpenin A5-sensitive succinate:decylubiquinone:DCIP reductase activity. Before measurements, cells were maintained for 48 hours in a medium containing glucose or galactose as the main carbon source, since galactose forces cells to use mitochondria to generate ATP (Manfredi *et al*, 1999). In fibroblasts, neither complex I nor II showed significant differences in activity between genotypes when tested in glucose or galactose media (Fig. 5A-D). In FA myofibroblasts, we observed significantly higher complex II activity in glucose (Fig. 5C), but not in galactose (Fig. 5D), whereas complex I activity was unchanged (Fig. 5A, B). Normal or increased activities of complexes I and II suggest that in FA cells FeS cluster levels are sufficient for the assembly of functional enzymes.

**Figure 5.**
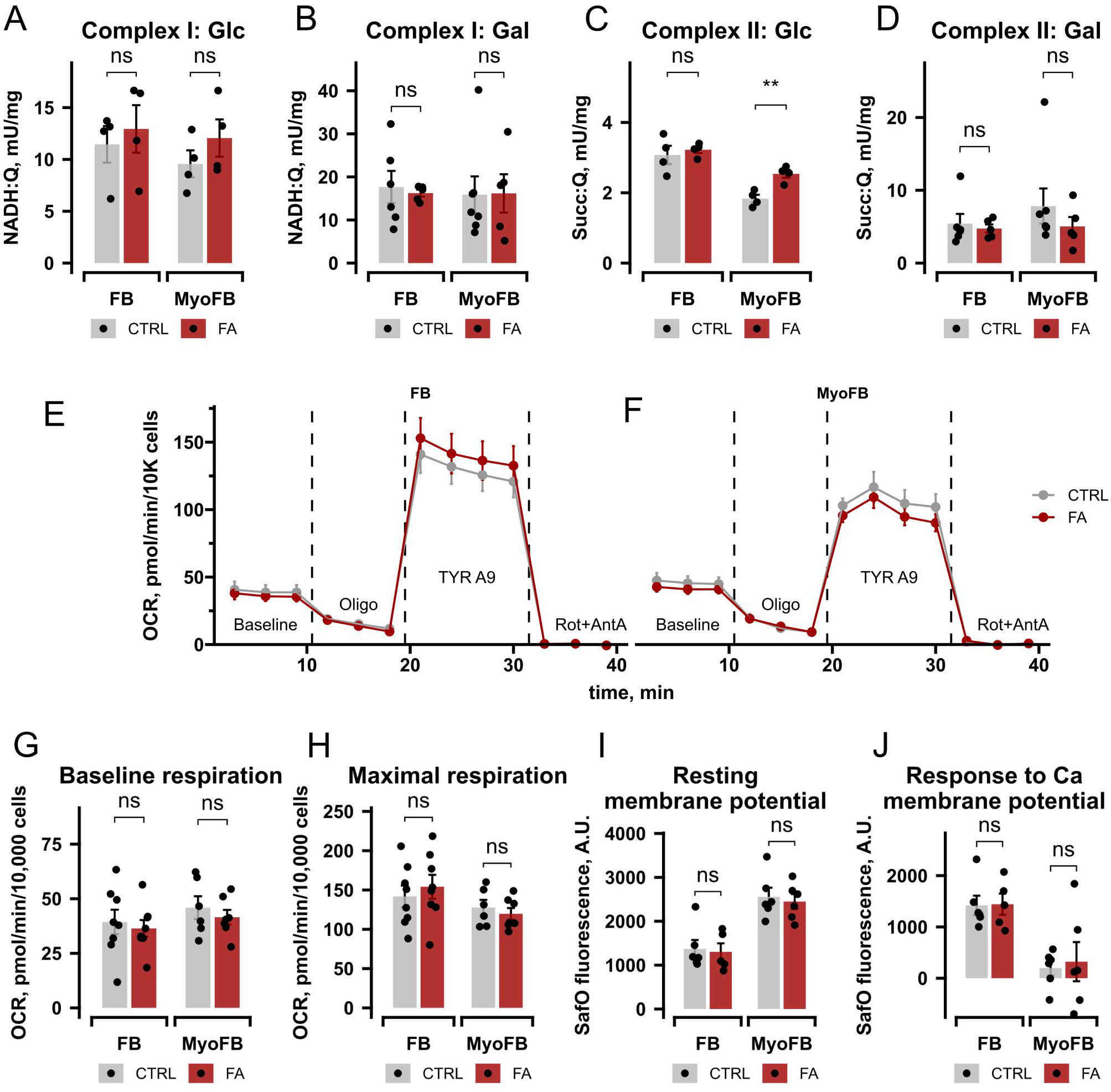
Mitochondrial bioenergetic function is preserved in FA fibroblasts and myofibroblasts. A, B. Complex I enzymatic activity in permeabilized fibroblasts and myofibroblasts maintained for 48 h in glucose (Glc, A) or galactose (Gal, B) media (n = 4-6 cell lines per group). Values are presented as rotenone-sensitive NADH:decylubiquinone reductase activity. C, D. Complex II enzymatic activity in permeabilized fibroblasts and myofibroblasts maintained for 48 h in glucose (C) or galactose (D) media (n = 4-6 cell lines per group). Values are presented as atpenin A5-sensitive succinate:decylubiquinone:DCIP reductase activity. E, F. Seahorse oxygen consumption rate (OCR) traces over time in fibroblasts (FB, E) and myofibroblasts (MyoFB, F) from control (CTRL, gray) and patient (FA, red) cell lines (n=6-8 cell lines per group); 4-5 technical replicates per plate from at least 2 independent plates were averaged for each cell line. Key injection points are indicated: Oligomycin (Oligo), tyrphostin A9 (TYR A9), and Rotenone/Antimycin A (Rot+AntA). G, H. Bar graphs of baseline OCR (G) and maximal OCR (H). Maximal respiration is defined as the maximum OCR measured following TYR A9 injection, minus the non-mitochondrial respiration after Rot/AntA (n=6-8 cell lines per group). I, J. Mitochondrial membrane potential represented by normalized Safranin O (SafO) fluorescence in arbitrary units (A.U.) in saponin-permeabilized fibroblasts and myofibroblasts (n = 5-6 cell lines per group) at baseline (I). Membrane potential changes in response to a 10µM Ca pulse (J). Mitochondria were energized with complex I substrates (glutamate, pyruvate, and malate). Lower fluorescence indicates more polarized mitochondria. Data information: All data represent mean ± SEM with individual data points overlaid. In E-H, Seahorse OCR data are normalized to cell count (10,000 cells). Statistical comparisons were performed using a two-tailed t-test: ns, not significant, * p < 0.05, ** p < 0.01.

Next, we measured cell respiration in control and FA fibroblasts (n=8/group) and myofibroblasts (n=6/group) grown in glucose medium. Baseline respiration, ATP-linked respiration upon addition of oligomycin, maximal respiratory capacity after injection of the uncoupler tyrphostin A9, and non-mitochondrial oxygen consumption rates (OCR) after addition of rotenone/antimycin A were assessed by Seahorse respirometry (Fig. 5E, F). We observed substantial line-to-line variability in both the control and FA groups for baseline and maximal OCR (Fig. 5G, H). Still, we found no significant differences between genotypes in either fibroblasts or myofibroblasts.

Finally, we measured mitochondrial membrane potential, the electrochemical driving force for both ATP synthesis and Ca uptake, using safranin O fluorescence in saponin-permeabilized fibroblasts and myofibroblasts (n=6). In this approach, lower fluorescence indicates a higher membrane potential because safranin accumulates in mitochondria and quenches fluorescence. Mitochondria were first energized with complex I substrates (glutamate, pyruvate, malate), then challenged with 10 µM Ca, followed by the addition of tyrphostin A9 to fully dissipate membrane potential. The resting membrane potential was not different between control and FA in fibroblasts or myofibroblasts (Fig. 5I). Upon Ca addition, membrane potential declined in fibroblasts, indicating that mitochondria took up Ca, and the decline was similar in control and FA (Fig. 5J). In contrast, Ca addition had no effect on membrane potential in myofibroblasts, confirming that their mitochondria do not take up significant amounts of Ca, as previously reported (Lombardi *et al*, 2019).

These bioenergetic measurements, spanning cellular respiration, membrane potential, and individual complex activities, demonstrate that, on average, mitochondrial function is preserved in FA fibroblasts and myofibroblasts despite FXN deficiency. Importantly, these results suggest that mitochondrial impairment is unlikely to be the mechanism underlying cytosolic Ca alterations in FA fibroblasts.

### *NCX1* expression normalizes cytosolic Ca and *CCN2* expression in FA cells

To investigate the molecular underpinnings of increased cytosolic Ca levels in FA fibroblasts, we mined our proteomics data for proteins involved in Ca homeostasis and signaling (Berridge *et al*, 2000). Of the 290 Ca-related proteins identified by proteomics, NCX1 (encoded by *SLC8A1*) was the most decreased one in FA fibroblasts (35% decrease, Fig. 6A). This finding was in accordance with the transcriptomics data showing decreased *SLC8A1* expression (Fig. 1A). Evaluation of known NCX1 transcriptional repressors or activators, including components of the inhibitory REST complex (Guida *et al*, 2024) or activating HIF1ɑ (Hudecova *et al*, 2011), by transcriptomics and proteomics analyses did not reveal significant differences between FA cells and controls (Appendix S2). Nevertheless, we found a significant direct correlation between *NCX1* and *FXN* gene expression (Fig. 6B). Since *FXN* levels inversely correlate with GAA repeat expansion length, our findings suggest that *FXN* levels or the GAA expansion itself can modulate *NCX1* expression epigenetically. Furthermore, we observed that baseline Ca levels inversely correlated with *NCX1* expression (Fig. 6C). Together, these findings suggest that decreased Ca extrusion from the cytosol through *NCX1* contributes to increased cytosolic Ca levels in FA fibroblasts.

**Figure 6.**
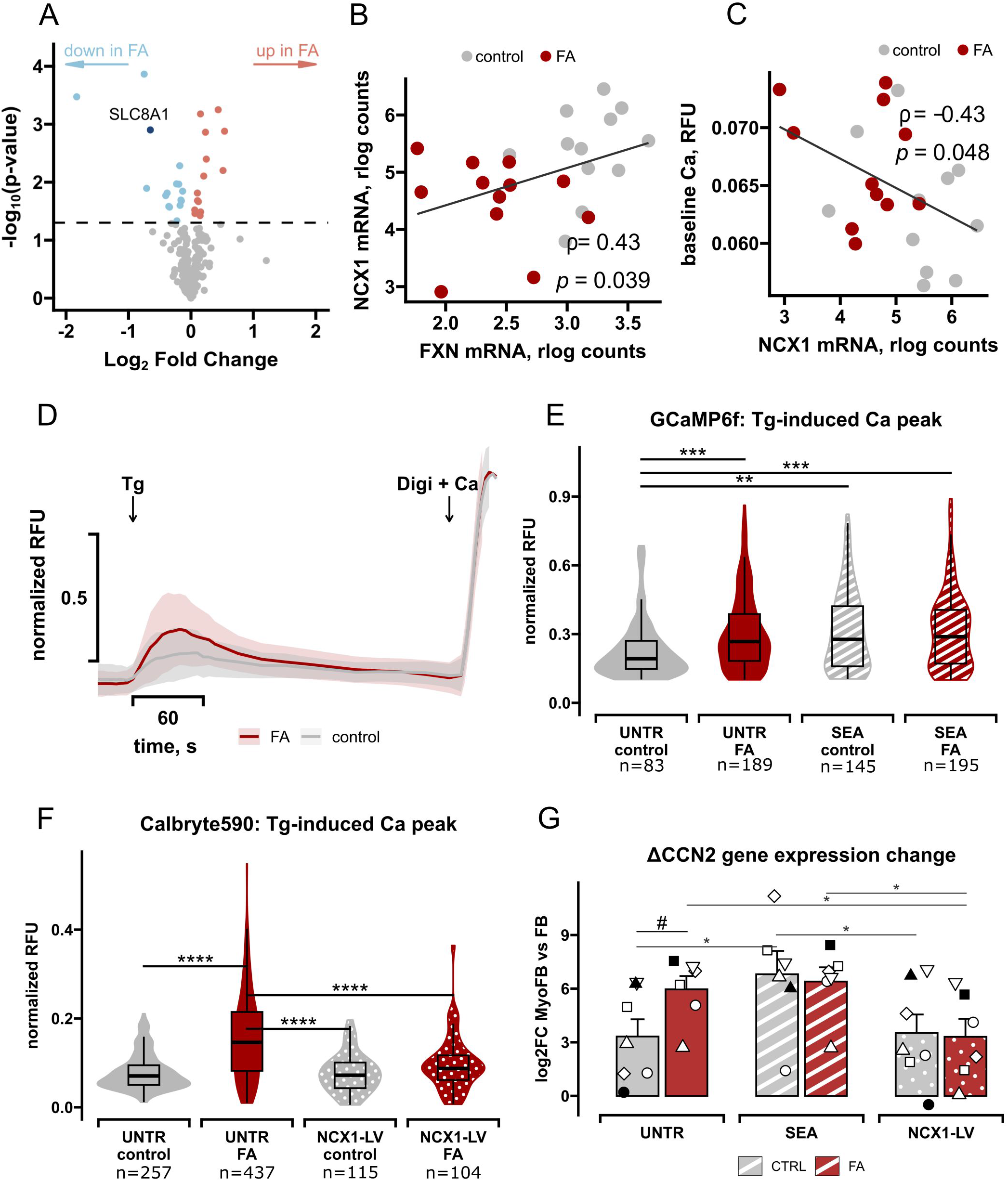
NCX1 modulation normalizes cytosolic Ca and CCN2 expression in FA cells. A. Volcano plot of Ca-related proteins from fibroblast proteomics. The negative log_10_ of p-value was plotted against log_2_FC, highlighting proteins with significantly different abundance between FA and control cells (p < 0.05). B, C. Spearman’s rank correlations between *FXN*, *NCX1*, and baseline Ca (n = 24 cell lines): *FXN* versus *NCX1* expression (B), baseline cytosolic Ca versus NCX1 expression (C). Regularized log (rlog) expression levels for FXN and NCX1 were used. Median values for each cell line for baseline Ca were used. D. Cytosolic Ca dynamics in response to thapsigargin (Tg, 1 µM) in fibroblasts at baseline, measured by GCaMP6f. Curves show median Ca traces constructed by calculating the median normalized RFU for each experimental group at every unified time point (from 6-8 cell lines per group). Arrows indicate the addition of Tg followed by digitonin (Digi, 10 μM) plus Ca (10 mM) for normalization. Shaded areas indicate variability represented by the median absolute deviation (MAD). E. Quantification of peak amplitude of Tg-induced Ca release (GCaMP6f fluorescence) in untreated fibroblasts in panel D (UNTR) and fibroblasts treated with the NCX1 inhibitor SEA0400 (SEA, 1 μM, 15 min). The number of cells measured is indicated for each group (from 6-8 cell lines per group). Violin plots show distribution density with internal boxplots indicating the median and interquartile range. F. Quantification of peak amplitude of Tg-induced Ca release measured with Calbryte 590, following lentiviral transduction of fibroblasts with NCX1-LV. The number of cells measured is indicated for each group (from 4 cell lines per group). Violin plots show distribution density with internal boxplots indicating the median and interquartile range. G. *CCN2* gene expression changes upon FMT. Log_2_ fold-change of *CCN2* mRNA expression between fibroblast and myofibroblast states (ΔCCN2 = CCN2_MyoFB_ - CCN2_FB_) in untreated (UNTR) cells or following chronic NCX1 inhibition with SEA (0.5 μM) for 4 days or transduction with NCX1-LV. Bars represent mean ± SEM, symbols indicate individual fibroblast lines subjected to different treatments. Data information: In B and C, data are analyzed by Spearman’s rank correlation; correlation coefficients ρ and associated p-values are shown. In E and F, asterisks indicate significance of Wilcoxon test: * p < 0.05, ** p < 0.01, *** p < 0.001. In G, to account for the dependency between treatments applied to the same cell lines, comparisons were performed using a linear mixed model with genotype and treatment as fixed factors and cell line as a random factor, with Estimated Marginal Means post-hoc tests (# p = 0.057, * p < 0.05).

Building on these findings, we investigated the role of NCX1 in FA Ca dysregulation. To induce Ca release from ER stores and prevent its reuptake by the ER, we used the sarco/endoplasmic reticulum Ca-ATPase (SERCA) inhibitor thapsigargin (Tg). We reasoned that this experimental paradigm forces cells to extrude Ca through the plasma membrane, a process dependent on NCX1, to maintain cytosolic Ca homeostasis. FA fibroblasts displayed significantly higher Tg-induced Ca peaks compared to control cells (Fig. 6D, E). This difference was more prominent than that observed with histamine (Fig. 4A, B), suggesting that defective plasma membrane extrusion contributes to elevated cytosolic Ca in FA fibroblasts.

To determine whether NCX1 loss drives this phenotype, we first pharmacologically blocked the exchanger with its specific inhibitor, SEA0400 (SEA) (Wang *et al*, 2007). In control fibroblasts, NCX1 inhibition was sufficient to recapitulate the FA phenotype, as evidenced by significantly elevated Tg-induced Ca peaks (Fig. 6D). On the other hand, SEA did not affect FA fibroblasts, suggesting that NCX1 activity was already low and could not be further inhibited in these cells.

Next, we tested the effects of restoring NCX1 levels in FA fibroblasts via lentiviral transduction. We transduced cells with a lentiviral construct (NCX1-LV) that co-expresses *EGFP* and human *NCX1* under independent promoters. Transduced cells were identified by green fluorescence (Appendix Fig. S3) and used for Ca measurements with Calbryte 590 (590nm emission) to avoid fluorescence spectral overlap with EGFP. The difference in Tg-induced Ca peak between untransduced control and FA fibroblasts was also confirmed using this approach (Fig. 6F). NCX1-LV did not affect Ca responses in control cells. In contrast, NCX1-LV rescued the FA phenotype, abolishing the differences in the Tg-induced Ca peak between FA and control.

Finally, we asked whether modulating NCX1 affects the fibrotic program by monitoring *CCN2* expression as a marker of fibrosis after treating fibroblasts with TGFβ to induce FMT. We calculated the fold change in *CCN2* expression during the FMT (*ΔCCN2*), as in Fig. 3D, in a subset of lines (7 controls and 6 FA) by qPCR. Consistent with transcriptomic data from the entire cohort (Fig. 3D), *CCN2* expression induction in myofibroblasts was higher in FA (Fig. 6G, UNTR). NCX1 inhibition by SEA exacerbated *CCN2* expression change in control lines (Fig. 6G, SEA), whereas NCX1 overexpression blunted the upregulation of *CCN2* in FA lines (Fig. 6G, NCX1-LV). These results mirrored the effects of SEA and NCX1-LV on Tg-induced Ca responses (Fig. 6E, F), thereby identifying decreased NCX1 as a cause of Ca alterations and profibrotic responses in FA fibroblasts.

## Discussion

The present study assessed whether human FA fibroblasts display intrinsic pro-fibrotic properties. The primary finding is that skin fibroblasts from FA patients are predisposed to pro-fibrotic reprogramming associated with defective cytosolic Ca clearance, in the absence of mitochondrial bioenergetic dysfunction. We identified NCX1 downregulation as the central mechanism underlying this cytosolic Ca dysregulation. Modulating NCX1, genetically or pharmacologically, in fibroblasts controlled the expression of pro-fibrotic factors, such as *CCN2*, during FMT. In control fibroblasts, chemical inhibition of NCX1 activity altered cytosolic Ca levels and enhanced *CCN2* expression, recapitulating FA phenotypes. In FA fibroblasts, NCX1 transduction restored cytosolic Ca dynamics and blunted *CCN2* expression in FMT.

We determined that cytosolic Ca dysregulation in FA cells is not the consequence of a mitochondrial bioenergetic failure. While mitochondrial dysfunction is a hallmark of FA in neural and cardiac tissues (Rötig *et al*, 1997; Bradley *et al*, 2000), we found that FA skin fibroblasts and myofibroblasts maintain normal mitochondrial respiratory chain function and membrane potential, which support the electrochemical gradient required for mitochondrial Ca uptake. Our findings highlight the challenge of identifying clear bioenergetic phenotypes in human FA fibroblasts. Previous literature shows no clear agreement on OXPHOS deficiency in human FA skin fibroblasts (Rötig *et al*, 1997; Abeti *et al*, 2018a), and FA myofibroblasts have never been investigated. We observed individual line-to-line variability in both FA and controls, which strongly advises including multiple biological replicates in OXPHOS studies. Moreover, fibroblasts are metabolically flexible and could adapt to limited FXN availability by relying on glycolysis instead of OXPHOS for ATP generation. The preservation of mitochondrial function in our cohort of FA fibroblasts and myofibroblasts suggests that these cellular models allow for the study of OXPHOS-unrelated FA mechanisms, such as fibrotic and Ca-handling properties.

Previous studies identified alterations in Ca homeostasis in different cell types from FA mouse models, including cardiomyocytes, cerebellar granule neurons (Abeti *et al*, 2018b), and astrocytes (Marullo *et al*, 2025). However, this is the first demonstration that cytosolic Ca is dysregulated in FA human fibroblasts. Earlier studies in mice have emphasized the importance of Ca regulation in fibroblast activation and highlighted the role of mitochondrial Ca regulation (Lombardi *et al*, 2019; Gibb *et al*, 2020). Although we do not exclude that mitochondrial Ca dynamics may also contribute to profibrotic phenotypes, our findings indicate that in FA fibroblasts plasma membrane extrusion via NCX1 plays a critical role. This interpretation is based on the downregulation of NCX1 and its genetic and pharmacological modulation in FA fibroblasts.

Downregulation of NCX1 expression has been reported in human transcriptomics datasets from FA skin fibroblasts (Napierala *et al*, 2017) and other cell types, such as iPSC-derived cardiomyocytes (Li *et al*, 2019). Here, we show that NCX1 expression levels directly correlate with FXN expression. FXN levels are known to inversely correlate with GAA repeat expansion (Lynch *et al*, 2024). Therefore, we speculate that FXN gene expansion negatively regulates NCX1 expression. Our transcriptomics and proteomics studies do not implicate the known inhibitory REST (Guida *et al*, 2024) or the activating HIF1ɑ (Hudecova *et al*, 2011) complexes in NCX1 downregulation in FA fibroblasts. This suggests two potentially overlapping alternative explanations.

First, FXN levels may modulate *NCX1* gene expression through yet unknown mechanisms. Speculatively, the FXN-NCX1 connection may involve transcription factors that require iron-sulfur clusters or iron-dependent zinc fingers, different from the previously reported MTF-1 (Valsecchi *et al*, 2021), which was not identified in our study among significant hits. However, transcriptomic data from mouse hearts where *FXN* was completely absent in cardiomyocytes (Payne *et al*, 2025), or extremely depleted due to an *FXN^G127V^* point mutation in the endogenous gene (Sayles *et al*, 2023), did not show *NCX1* downregulation. These findings suggest that FXN protein levels do not regulate *NCX1* expression in mice. However, human and rodent *NCX1* gene regulatory regions differ, as shown by sequence alignment in the Genome Browser (hg38,chr2:40,097,270-40,452,090) (Casper *et al*, 2026). Moreover, in human iPSC-derived cardiomyocytes in which *FXN* was acutely silenced for one week, the expression of *NCX1* was not significantly altered (Cotticelli *et al*, 2023). Taken together, this evidence suggests that FXN loss alone is insufficient to cause NCX1 downregulation.

Second, *NCX1* repression could result from GAA-repeat silencing, since DNA repeat expansions, including those in the *FXN* gene, have been shown to be associated with chromatin remodeling and heterochromatin formation (Kumari & Usdin, 2009). However, in FA human fibroblasts, the GAA expansion was shown to affect only *FXN* expression, not other genes on chromosome 9 (Li *et al*, 2015). Therefore, whether genes on other chromosomes, including *NCX1* on chromosome 2, could be affected by chromatin remodeling caused by GAA repeats remains to be determined. Future work, including epigenetic regulation of *NCX1* expression, will investigate how *NCX1* is transcriptionally repressed in human FA fibroblasts.

The effects of NCX1 up- or downregulation are highly tissue- and disease-specific and may relate to whether NCX1 operates in forward mode (Ca extrusion) or reverse mode (Ca uptake). In the heart, NCX1 upregulation was linked to increased risk of cardiac hypertrophy and arrhythmias, and its pharmacological or genetic inhibition was proposed to be protective (Hobai & O’Rourke, 2004). In contrast, in the brain, NCX1 upregulation contributed to the neuroprotective effect of ischemic preconditioning by limiting ER Ca load (Formisano *et al*, 2013), while NCX1 inhibition worsened ischemic outcomes (Valsecchi *et al*, 2021). In the kidney, loss of NCX1 triggered profibrotic transformation of epithelial cells into fibroblasts (Balasubramaniam *et al*, 2017), suggesting that NCX1 dysregulation can play a pathogenic role in tissue fibrosis. However, the impact of NCX1 downregulation in cardiac fibrosis has not been investigated.

Although this study defined the profibrotic role of NCX1 inhibition and downregulation using human FA skin fibroblasts, validating this mechanism in cardiac fibrosis will require FA heart models. Skin and cardiac fibroblasts share common fibroblast cell markers but differ in tissue-specific markers (Zhang *et al*, 2019). Moreover, fibroblast monocultures cannot recapitulate the tissue microenvironment. To validate the hypothesis that NCX1 downregulation drives pathological fibrosis in the FA heart, human cardiac fibroblasts could be investigated using heart biopsies or iPSC-derived organoids containing all major cardiac cell types, including fibroblasts. Therefore, while our study identified cell-autonomous alterations focused on profibrotic transcriptional responses in fibroblasts, it could not fully address fibrotic events that occur *in vivo*. Despite these limitations, our results suggest that human FA skin fibroblasts could serve as a platform for small-molecule screening of NCX1 activators to attenuate profibrotic responses. Research on NCX1 activators remains in its early stages relative to the broad panel of available inhibitors (Scognamiglio *et al*, 2025). Moreover, whether alterations in *NCX1* expression in other disease-relevant cell types, including cardiomyocytes, have beneficial or detrimental effects needs to be determined to decide whether to target NCX1 broadly or specifically in fibroblasts to avoid adverse effects.

In conclusion, our study suggests that cytosolic Ca regulation modifies profibrotic stimulation in FA. We propose the following mechanism for intrinsic profibrotic bias in human FA fibroblasts: *NCX1* downregulation contributes to slower cytosolic Ca clearance, thereby sustaining Ca-dependent signaling and consequently increased expression of *CCN2* and other profibrotic factors. If validated in FA heart, our findings would identify NCX1-mediated Ca extrusion in fibroblasts as a potential therapeutic target to mitigate fibrosis that contributes to lethal cardiomyopathy in FA patients.

## Methods

### Human Fibroblast Cell Lines

All anonymized fibroblast cell lines were obtained from the Friedreich’s Ataxia (FA) cell repository in Dr. Napierala’s laboratory at UT Southwestern Medical Center. Characterization of GAA repeat length in the FXN gene was previously described (Li *et al*, 2016). Passage numbers at the time of experiments ranged from p5 to p15. We used patient demographics, including age, sex, and GAA repeat lengths, for correlational analyses. Fibroblasts were cultivated in complete Dulbecco’s Modified Eagle’s Medium (DMEM, Gibco, 11995065) supplemented with 10% fetal bovine serum (FBS), 1% antibiotic/antimycotic, and plasmocin (2.5 µg/mL, InvivoGen). All experiments were conducted within 3 passages of thawing. We used 12 FA patient-derived and 12 age- and sex-matched control fibroblast lines for transcriptomic profiling. Randomized subsets of these cell lines were used for proteomic, Ca imaging, bioenergetic and NCX1 modulation experiments. Randomization was performed on the FA group and sex-age matching controls were selected for each random FA subset of lines. Cells were analyzed in two states, fibroblasts (FB) and TGFβ1-induced myofibroblasts (MyoFB).

Myofibroblast differentiation was induced as previously described (Piersma *et al*, 2017) with minor modifications. Briefly, fibroblasts were seeded at a density of 8,000 cells/cm² in complete DMEM on fibronectin-coated plates (2 µg/cm²). After 24 hours, serum starvation was initiated by replacing the growth medium with DMEM supplemented with 0.5% FBS for 16-18 hours. Myofibroblast differentiation was then induced by treatment with 10 ng/mL recombinant human TGFβ1 (R&D Systems, 7754-BH-025) in DMEM supplemented with 1% FBS for 72 hours (transcriptomics, proteomics, qPCR) or up to 5 days (immunocytochemistry, Western blot). Medium containing TGFβ1 was replaced every 48 hours. Successful differentiation was confirmed by upregulation of αSMA and Collagen I, as assessed by Western blot and immunocytochemistry.

### Lentiviral Transduction

For NCX1 overexpression, fibroblasts at 60–70% confluence were transduced with the pLV[Exp]-EGFP/Puro-EF1A-hSLC8A1 lentiviral vector (VectorBuilder ID: VB900167-3938rak) encoding human SLC8A1 (NCX1, NM_001394103.1) at a multiplicity of infection (MOI) of 10 in the presence of 8 µg/mL polybrene in the antibiotic-free experimental medium DMEM (A14430), containing 5 mM glucose, 10% FBS, pyruvate and GlutaMAX. After 24 hours, the transduction medium was removed and replaced with experimental medium. Cells were cultured for an additional 72 hours prior to downstream analysis. Transduction efficiency was assessed by EGFP fluorescence using fluorescence microscopy.

For live-cell Ca imaging, fibroblasts were transduced with an LV-CAG-GCaMP6f lentiviral vector (SignaGen, SL100321) at an MOI of 10 in the presence of 6 µg/mL polybrene in the antibiotic-free experimental medium. After 24 hours, the medium was replaced with fresh experimental medium, and cells were allowed to recover for 48 hours before imaging.

### Pharmacological Treatments

For chronic NCX1 inhibition, fibroblasts were treated with 0.5 µM SEA0400 (Tocris Bioscience, 6140, 50 mM in DMSO) for 4 days, replenishing at each medium change. Vehicle-treated cells served as controls.

For acute NCX1 inhibition in Ca imaging experiments, SEA0400 was applied at 1 µM for 15 minutes immediately prior to imaging.

### 3’-RNA Sequencing

Total RNA was extracted from fibroblasts (FB) and TGFβ1-induced myofibroblasts (MyoFB) from 12 FA and 12 control cell lines using the TRIzol/chloroform method. Cells from 6-cm dishes were washed with ice-cold PBS and lysed in 0.5 mL TRIzol; 0.1 mL chloroform was added, and RNA was purified from the aqueous phase using the SV Total

RNA Isolation System (Promega, A3800). RNA integrity was confirmed by agarose gel electrophoresis and NanoDrop spectrophotometry (A260/A280>2.0 and A260/A230>1.6). 3′ mRNA library preparation and sequencing were performed by the Cornell Genomics Core on an Illumina NextSeq 500 (Flowcell: HTKJMBGXN) with a High Output 75-cycle kit configured for 86 bp single-end reads.

Raw reads were quality-filtered with fastp (v0.20.1) (Chen *et al*, 2018) and quantified with Salmon (v1.10.2) (Patro *et al*, 2017) against a custom human “gentrome” reference index built from GRCh38 transcripts and long non-coding RNA with primary genomic assembly as decoy (GENECODE Release 49, GRCh38.p14 (Nurk *et al*, 2022)). Quantification included sequence-bias, positional-bias, and GC-content bias corrections (--seqBias, --posBias, --gcBias) with selective alignment (--validateMappings). All downstream analysis was conducted in RStudio (v4.5.1+) using Bioconductor packages. Transcript-level counts were imported and aggregated to the gene level using *tximport*. Only protein-coding and lncRNA biotypes (filtered via biomaRt package) were used for downstream analysis.

#### Differential expression analysis

Differential expression analysis was performed using *DESeq2* (v1.50.2) (Love *et al*, 2014). Two different study designs were used, one-way and two-way. To assess differences between FA and CTRL within each type, we calculated independent contrasts: FA_FB vs CTRL_FB or FA_MyoFB vs CTRL_MyoFB, with a design *∼ group*, where each group factor was a combination of the original genotype and cell type factors. To assess differences in TGFβ1 response between FA and CTRL during fibroblast-to-myofibroblast transition, we investigated the interaction term from a design *∼ cell type + genotype + genotype:cell type*. Genes with p_adj_-value < 0.05 (Benjamini-Hochberg correction) were considered differentially expressed. Regularized log (rlog) counts were used for visualization and correlation analyses. **ΔGene expression calculation.** The transcriptional response to TGFβ1 per cell line was quantified as: *ΔGene = rlog(Gene_MyoFB) − rlog(Gene_FB)*, where rlog values were computed jointly across all 48 samples. This metric was used for correlation with FXN expression.

#### Weighted Gene Co-Expression Analysis (WGCNA)

Co-expression network analysis used the *WGCNA* (v1.74) and *flashClust* (1.1-4) R packages and variance-stabilized normalized count data from DESeq2. A soft-thresholding power of 4 was selected via *pickSoftThreshold* function. A signed co-expression network was constructed with a minimum module size of 30 genes. Module eigengenes, representing the first principal component of module expression profiles, were correlated with sample traits (disease status, GAA repeat length, FXN mRNA expression, age, sex) using Spearman’s rank correlation; modules with p_adj_-value < 0.05 were considered trait-associated.

### Gene Set Enrichment Analysis (GSEA) and pathway enrichment

For FB-to-MyoFB transition analysis, GSEA was performed using the *clusterProfiler* (v4.18.4) package (Wu *et al*, 2021). Genes were ranked by their DESeq2 Wald statistic to incorporate both the direction and statistical strength of the changes. Ranked gene lists were queried against the KEGG and Reactome databases, restricting the analysis to sets of 10 to 500 genes per term and FDR-correcting throughout (p adj < 0.05). For WGCNA module composition analysis, unranked module genes for each cluster were compared against Gene Ontology (GO) Biological Process database, and redundant terms were reduced using semantic similarity clustering (cutoff =0.7) to merge redundant GO terms.

#### Transcription factor (TF) activity

For each sample, TF activity scores were inferred from normalized rlog counts using *decoupleR* (v2.16) (Badia-I-Mompel *et al*, 2022), which integrates gene transcription levels with a curated TF-gene interaction network from CollectTRI. We used univariate linear modeling, multivariate linear modeling, and normalized weighted sum algorithms, followed by consensus score estimation that integrated all methods. TF activity scores were compared between FA and CTRL samples using a t-test. TFs with p_adj_-value < 0.05 were correlated with FXN mRNA expression using Spearman’s rank correlation. *ComplexHeatmap* (v2.26.1) package was used for visualization (Gu *et al*, 2016).

### Fibrotic and Ca Gene Panel Curation

A consensus panel of fibrotic genes was assembled via systematic literature-driven text mining using Cytoscape (stringApp v2.2.0) (Doncheva *et al*, 2019). PubMed queries used the following search terms: (’fibrosis’ OR ‘profibrotic’ OR ‘myofibroblast’) AND (’TGFβ signaling’ OR ‘extracellular matrix’ OR ‘ECM remodeling’) AND (’marker’ OR ‘signature’ OR ‘gene expression’), restricted to 250 entries. STRING interaction confidence scores were used for prioritization, and tissue expression data were filtered for fibroblast-intrinsic candidates. The final panel of genes with established roles in TGFβ1 signaling, ECM remodeling, and myofibroblast transition is provided in (Dataset EV1).

A list of proteins associated with Ca signaling and dynamics was constructed following a previously reported approach (Hörtenhuber *et al*, 2017) with modifications. The foundation was built based on the known signal transduction elements, receptors, channels and downstream effectors outlined in (Berridge *et al*, 2000) and expanded using *g:Profiler* (Reimand *et al*, 2019) by integrating Ca-associated GO terms and KEGG pathways, such as Calcium Channel Complex (GO:0034704), Cellular Calcium Ion Homeostasis (GO:0006874), Response to Calcium Ion (GO:0051592), and the cAMP Signaling Pathway (KEGG:04024). The curated list of proteins involved in Ca regulation and Ca-target proteins responsive to intracellular Ca fluxes is provided in (Dataset EV2).

### Proteomics and Phosphoproteomics

Fibroblasts from 8 control and 8 FA cell lines were grown in 10-cm dishes, washed twice with ice-cold PBS, and lysed by scraping in 300 µL of freshly prepared 9 M urea buffer (50 mM Tris-HCl pH 8.0, 5 mM NaF, 1 mM Na₃VO₄, supplemented with cOmplete protease inhibitor cocktail (Roche)). Lysates were stored at -80°C until submission to the Weill Cornell Medicine Proteomics and Metabolomics Core Facility. Proteins were precipitated and digested with trypsin. Desalted peptides were labeled by 16-plex TMT. For phosphoproteomics, we used≥ 1 mg total protein per condition; for expression profiling, ≥ 100 µg per condition. Phosphopeptides were enriched by TiO₂. Raw MS data were acquired on Orbitrap Fusion Lumos and searched against the UniProt human proteome (UP000005640) using MaxQuant (v2.3+) at 1% FDR at both protein and peptide levels. A total of 9,128 proteins were identified (7,899 quantified), and 23,856 phosphosites were identified (18,266 quantified). Protein and phosphopeptide intensities were log₂-transformed and median-normalized per sample. Differential abundance was assessed by a two-sample t-test with Benjamini-Hochberg FDR correction (p_adj_-value < 0.05). Phosphosites were mapped to canonical UniProt sequences and classified as serine, threonine, or tyrosine phosphorylations. Omics Vizualizer (v1.3.1) was used within Cytoscape environment to depict protein fold change and phosphopeptide fold change simultaneously (Legeay et al, 2020).

### Live-Cell Ca Imaging

Cells were seeded on a 96-well glass-bottom black plate (Cellvis, P96-1.5H-N) at 8,000 cells/cm² and allowed to adhere for 24 hours prior to lentiviral transduction with LV-CAG-GCaMP6f or Calbryte 590 AM dye loading. Cells were maintained in Krebs-Ringer modified buffer (KRB, 135 mM NaCl, 5 mM KCl, 1 mM MgSO4, 1 mM MgCl2, 0.4 mM KH2PO4, 20 mM HEPES, 1 mM CaCl2, pH 7.4) at 25°C throughout imaging. For Ca imaging after NCX1 overexpression, fibroblasts were loaded with 5 µM Calbryte 590 AM (ATT Bioquest, 20700, 5 mM in DMSO) in KRB buffer containing 0.04% Pluronic F-127 for 60 min at 37°C, followed by a 30-min washing period.

Imaging was performed using a Leica DM IRB inverted fluorescence microscope equipped with a Leica N PLAN L 20× objective lens (NA 0.40, ∞/0–2, LMC; Leica, #506134), CoolLED pE-300 LED light source, filter sets, and a 12-bit Retiga 1350B monochrome CCD camera (QImaging) with 2×2 binning. Image acquisition was controlled via Micro-Manager software (v2.0.0) (Edelstein et al, 2014).

GCaMP6f fluorescence was excited for 250 ms using the blue light channel (100% intensity) and collected every 0.8-2 s with Chroma 59001v2 ET - DAPI/Green FISH filter set (excitation 379-498 nm and 470-495 nm, emission 425-460 nm and 500-545 nm).

Calbryte 590 AM fluorescence was excited for 100 ms using the green light channel (100% intensity) and collected every 2 s with the Leica N2.1 filter set (excitation 515-560 nm, long-pass emission >590 nm).

After recording a 30-second baseline, cytosolic Ca transients were evoked by adding histamine (100 µM final) or thapsigargin (1 µM), and cells were monitored for 3-5 minutes. Digitonin (10 µM) and Ca (10 mM) were subsequently added to achieve maximal fluorescence.

Cell segmentation and fluorescence (mean gray values) extraction were performed in ImageJ Fiji (v1.53+). Individual cell fluorescence traces were aligned to the rise time of the histamine/thapsigargin-induced response, resampled onto a unified time grid using local regression (LOCFIT), and normalized to the maximal digitonin/Ca signal to calculate a relative fluorescence unit (RFU) ratio. Peak amplitude, baseline fluorescence, area under the curve (AUC), and decay rate (k_decay) were quantified using custom R scripts in RStudio.

### Mitochondrial Assays

#### Oxygen Consumption Rate

Mitochondrial respiration was assessed using a Seahorse XF96 Extracellular Flux Analyzer (Agilent). Fibroblasts were seeded at 10,000 cells/well and myofibroblasts at 3,000 cells/well (prior to TGFβ1 treatment). On assay day, growth medium was replaced with 200 µL Agilent Seahorse XF Base Medium (Agilent, 103334-100) supplemented with 2 mM L-glutamine (Gibco, 25030-081), 1 mM sodium pyruvate (Gibco, 11360-070), and 5 mM D-glucose (Sigma, G8769), and cells were incubated for 1 hour in a non-CO₂ incubator at 37°C. Compounds were loaded into injection ports and delivered sequentially: oligomycin (0.5 µg/mL); uncoupler tyrphostin A9 (0.5 µM); rotenone (0.5 µM) + antimycin A (1 mM). Raw OCR was normalized to the final cell count per well (pmol/min per 10,000 cells) determined by Hoechst staining and automated counting on ImageXpress Pico (Molecular Devices). Non-mitochondrial respiration (after Rot/AntA) was subtracted from all values.

#### Mitochondrial Membrane Potential

Mitochondrial membrane potential (ΔΨm) was assessed in permeabilized cells using Safranin O fluorescence as described (Figueira *et al*, 2012) with modifications. Fibroblasts and myofibroblasts were harvested using Accutase (Innovative Cell Technologies, AM105) and counted. For each assay, 200,000 cells were resuspended in 200 µL permeabilization buffer (8 mM KCl, 100 mM potassium gluconate, 10 mM NaCl, 10 mM HEPES, 10 mM KH₂PO₄, 0.5 µM EGTA, 10 mM mannitol, 1 mM MgCl₂, 0.5 mg/mL BSA, pH 7.2) containing saponin (50 µg/mL) and permeabilized for 5-10 minutes at room temperature. Fluorescence was recorded on a SpectraMax M5 plate reader (Molecular Devices; excitation 495 nm, emission 587 nm) using white opaque 96-well plates at 37°C. Mitochondrial substrates (5 mM glutamate, 5 mM pyruvate, 2 mM malate) were added, followed by Safranin O (5 µM). Sequential additions of Ca (10 µM) and tyrphostin A9 (1 µM) were used to assess Ca-induced changes in membrane potential and full mitochondrial depolarization, respectively. Normalized ΔΨm was calculated as: *ΔΨm_norm = (F_max − F_rest) / (F_max − F_min)*, where F_max is fluorescence after full uncoupling, F_rest is resting fluorescence, and F_min is the minimum fluorescence value.

#### Mitochondrial Complex I and II Activities

Complex I (NADH:decylubiquinone reductase) and Complex II (succinate:DBQ:DCIP reductase) activities were measured spectrophotometrically in mildly permeabilized cells using the SpectraMax M5 plate reader (Molecular Devices) at 25°C.

Cells from a confluent well of a 6-well plate were collected in the isolation buffer (10 mM Tris-HCl pH 7.5, 1 mM EGTA, 1 mM EDTA, 225 mM mannitol, 75 mM sucrose) and pelleted. Cell pellets were lysed in 20 mM HEPES-KOH (pH 7.8) containing 0.025% n-dodecyl maltoside and 1 mM citrate. Complex I activity was measured in 200 µL reaction buffer (20 mM HEPES pH 7.8, 30 µg/mL alamethicin (Cayman Chemical) supplemented with 1 mg/mL BSA and 1 mM KCN, using 0.15 mM NADH as electron donor and 40 µM decylubiquinone (DBQ) as acceptor with 5-15 µg protein/well. NADH oxidation was monitored at 340 nm (ε = 6.22 mM⁻¹cm⁻¹); rotenone-sensitive activity was used to define Complex I-specific activity.

Complex II activity was assayed under identical buffer conditions using 10 mM succinate, 50 µM DBQ, and 50 µM DCIP with 2-5 µg protein/well, monitored at 600 nm (ε = 21 mM⁻¹cm⁻¹); atpenin A5-sensitive activity defined Complex II-specific signal.

### Western Blotting

Cells were lysed in RIPA buffer (20 mM Tris-HCl pH 7.4, 150 mM NaCl, 1% Triton X-100, 0.1% SDS, 1% sodium deoxycholate, 1 mM EDTA) supplemented with cOmplete protease inhibitor cocktail (Roche). Protein concentration was determined by BCA assay (ThermoFisher). Fifteen micrograms of total protein per lane were resolved on AnyKD gradient SDS-PAGE gels (BioRad) at 70V for 10 min followed by 200V until completion. Proteins were transferred to nitrocellulose membranes using a Trans-Blot Turbo System (BioRad) at 2.5A for 15 minutes. Membranes were blocked with 5% non-fat dry milk in TBS containing 1% Tween-20 (TBST) for 30-60 minutes at room temperature and incubated overnight at 4°C with primary antibodies. After washing in TBST, membranes were incubated with HRP-conjugated secondary antibodies, anti-rabbit (111-035-144, Jackson ImmunoResearch, 1:10,000) or anti-mouse (115-035-146, Jackson ImmunoResearch, 1:10,000) for 30 minutes at room temperature. Chemiluminescence was developed using Clarity Western ECL substrate (BioRad). Images were processed using ImageLab software (BioRad). For primary antibodies incubations, the same membrane underwent sequential probing and stripping in the following order. First, rabbit anti-collagen I (Col I, Abcam, ab260043, 1:1000, 4°C, overnight), subsequently anti-vimentin (Vim, GeneTex, GTX100619, 1:50,000, room temperature, 1 h). The membrane was then stripped, re-blocked, and probed for mouse anti-ɑSMA (αSMA, ThermoFisher, 14-9760-82, 1:1500, room temperature, 1 h). Finally, the membrane underwent a second strip-and-block cycle before probing for mouse anti-GAPDH (Abcam, ab8245, 1:10,000, room temperature, 1 h).

### Immunocytochemistry

Cells were seeded on 12mm round cover glass and fixed with 3% paraformaldehyde in PBS-Gly (20 mM Glycine in PBS) for 15 minutes at room temperature. Cells were permeabilized with 0.3% Triton X-100 in PBS-Gly for 7 minutes and blocked with 1% BSA and 0.1% Triton X-100 in PBS-Gly for 30 minutes. Primary antibodies against αSMA (ThermoFisher, 14-9760-82, 1:100) and Vimentin (GeneTex, GTX100619, 1:200) were applied in blocking buffer overnight at 4°C. Following 3 × 5 min PBS-Gly washes, cells were incubated with Alexa Fluor-conjugated secondary antibodies, Alexa 555 goat anti-mouse (Invitrogen, A21426, 1:1000) and Alexa 488 goat anti-rabbit (Invitrogen, A32731, 1:1000), for 30 minutes at room temperature in the dark. Nuclei were counterstained with Hoechst (5 µg/ml, Sigma). Coverslips were mounted using Fluoromount-G™ Mounting Medium. Images were captured using a Leica DM IRB inverted fluorescence microscope equipped with a Leica N PLAN L 40× objective lens (NA 0.55, CORR; Leica, #506297), without binning. DAPI/Hoechst and Green/FITC channels were captured using a Chroma 59001v2 ET - DAPI/Green FISH filter set (excitation 379–398 nm and 470–495 nm, emission 425–460 nm and 500–545 nm) with UV light excitation (200 ms exposure, 30% intensity) and blue light excitation (200 ms exposure, 20% intensity), respectively. Alternatively, mCherry channel was captured using a Chroma 59022 ET - EGFP/mCherry dual-band filter set (excitation 450–490 nm and 550–590 nm, emission 500–540 nm and 600–660 nm) with green light excitation (200 ms exposure, 40% intensity).

### Quantitative PCR (qPCR)

Total RNA was isolated using the Promega SV Total RNA Isolation System (Promega, Z3100) per manufacturer’s instructions. RNA concentration and purity were assessed by NanoDrop spectrophotometry. Complementary DNA (cDNA) was synthesized from ∼1,000 ng total RNA using the Improm-II Reverse Transcription System (Promega, A3800) with Oligo(dT) primers per manufacturer’s protocol. cDNA was diluted 5-fold prior to qPCR amplification. Reactions were performed on a QuantStudio 6 Flex System thermocycler using Applied Biosystems SYBR Green PCR Master Mix (ThermoFisher, 4309155) in a final volume of 20 µL. The reference gene β-actin (ACTB) was assayed using the QIAGEN QuantiTech Primer Assay (QT00095431). CCN2 (NM_001901.4) was amplified using custom primers: forward 5′-AAAAGTGCATCCGTACTCCCA-3′; reverse 5′-CCGTCGGTACATACTCCACAG-3′. Relative expression was calculated using the 2^−ΔΔCT^ method, normalized to ACTB.

### Quantification and Statistical Analysis

All statistical analyses were performed in RStudio (Posit team, 2026) using custom RMarkdown scripts (R version 4.4+). Data are presented as mean ± SEM for normally distributed data or as violin plots with box blots indicating mean and interquartile range for non-normally distributed data. The number of biological replicates (individual cell lines, n) for each experiment is specified in the corresponding figure legend. Technical replicates were averaged prior to statistical analysis. Normality of data distributions was assessed using the Anderson-Darling test. Normally distributed data were analyzed by an unpaired two-sample t-test. Non-normally distributed data were analyzed by Wilcoxon rank-sum test or Kruskal-Wallis test. Spearman’s rank correlation (ρ) was used for all correlation analyses. For NCX1 modulation experiments in which multiple treatments were applied to the same cell lines, a linear mixed-effects model was used with genotype and treatment as fixed effects and cell line as a random effect, with post-hoc comparisons using Estimated Marginal Means (emmeans R package, v2.0.3). Multiple comparison correction was performed using the Benjamini-Hochberg FDR method. A significance threshold of p < 0.05 was applied.

## Data and Code Availability

Raw RNA-seq data (FASTQ files) and processed count matrices have been deposited in the NCBI Gene Expression Omnibus (GEO) under accession number (to be assigned upon submission). Raw mass spectrometry data have been deposited in (to be assigned upon submission). All other data and custom R scripts supporting the findings are available upon request.

## Author Contribution

A.S., H.K., and G.M. contributed to conceptualization. A.S. was responsible for methodology, developing custom scripts in R, formal analysis, validation, investigation, data curation and visualization. A.S. and G.M. provided resources, supervised the study, and acquired funding. A.S., H.K., and G.M. managed project administration. A.S. and G.M. wrote the original draft. A.S., H.K., and G.M. reviewed and edited the manuscript.

## Disclosure and competing interest statement

The authors declare that they do not have any competing interests.

## Acknowledgements

We thank Ms. Jenipher Tenesaca and Ms. Grace Mao for their contribution to the experiments during their high-school internships. We thank Dr. Jordi Magrané for critically reading the manuscript, and Dr. Veronica Granatiero for setting up the cell culture system.

## Funding

This project was supported by grants from Friedreich’s Ataxia Research Alliance (FARA) to G.M. and A.S. and from National Institute of Health (NIH) R35 NS122209 to G.M.

## Expanded View Figure Legends

**Figure EV1. Concordant genes in FA fibroblasts and myofibroblasts**

A. UpSet plot showing differentially expressed genes in fibroblasts (FB) and myofibroblasts (MyoFB) and intersections between gene lists. Grey bars represent total upregulated or downregulated gene sets. Blue bar shows genes concordantly downregulated in both FA fibroblasts (FA FB Down) and myofibroblasts (FA MyoFB Down; n = 11 genes). Red bar shows genes concordantly upregulated in FA cells (FA FB Up and FA MyoFB UP; n = 9 genes).

B. Heatmap of log_2_ fold-change (log_2_FC) values for the 20 concordantly regulated genes across FB and MyoFB. Color gradient shows upregulation (red) and downregulation (blue) relative to control cells, with values indicated for each gene.

Data information: Gene set lists were obtained from differential gene expression analysis comparing FA versus control cells, for fibroblasts and myofibroblasts separately. Genes that passed the p_adj_-value threshold of 0.05 were considered significantly down- or upregulated.

### Dataset description

**Dataset EV1. Consensus panel of fibrotic genes assembled via systematic literature-driven text mining using Cytoscape**

**Dataset EV2. Curated list of proteins involved in Ca regulation and Ca-target proteins responsive to intracellular Ca fluxes**

## References

Abeti R, Baccaro A, Esteras N & Giunti P (2018a) Novel Nrf2-Inducer Prevents Mitochondrial Defects and Oxidative Stress in Friedreich’s Ataxia Models. Front Cell Neurosci 12: 188

Abeti R, Brown AF, Maiolino M, Patel S & Giunti P (2018b) Calcium Deregulation: Novel Insights to Understand Friedreich’s Ataxia Pathophysiology. Front Cell Neurosci 12: 264

Adapala RK, Thoppil RJ, Luther DJ, Paruchuri S, Meszaros JG, Chilian WM & Thodeti CK (2013) TRPV4 channels mediate cardiac fibroblast differentiation by integrating mechanical and soluble signals. J Mol Cell Cardiol 54: 45–52

Badia-I-Mompel P, Vélez Santiago J, Braunger J, Geiss C, Dimitrov D, Müller-Dott S, Taus P, Dugourd A, Holland CH, Ramirez Flores RO, et al (2022) decoupleR: ensemble of computational methods to infer biological activities from omics data. Bioinform Adv 2: vbac016

Balasubramaniam SL, Gopalakrishnapillai A, Petrelli NJ & Barwe SP (2017) Knockdown of sodium-calcium exchanger 1 induces epithelial-to-mesenchymal transition in kidney epithelial cells. J Biol Chem 292: 11388–11399

Baum J & Duffy HS (2011) Fibroblasts and myofibroblasts: what are we talking about? J Cardiovasc Pharmacol 57: 376–379

Berridge MJ, Lipp P & Bootman MD (2000) The versatility and universality of calcium signalling. Nat Rev Mol Cell Biol 1: 11–21

Bradley JL, Blake JC, Chamberlain S, Thomas PK, Cooper JM & Schapira AH (2000) Clinical, biochemical and molecular genetic correlations in Friedreich’s ataxia. Hum Mol Genet 9: 275–282

Campuzano V, Montermini L, Moltò MD, Pianese L, Cossée M, Cavalcanti F, Monros E, Rodius F, Duclos F, Monticelli A, et al (1996) Friedreich’s ataxia: autosomal recessive disease caused by an intronic GAA triplet repeat expansion. Science 271: 1423–1427

Casper J, Speir ML, Raney BJ, Perez G, Nassar LR, Lee CM, Hinrichs AS, Gonzalez JN, Fischer C, Diekhans M, et al (2026) The UCSC Genome Browser database: 2026 update. Nucleic Acids Res 54: D1331–D1335

Chen S, Zhou Y, Chen Y & Gu J (2018) fastp: an ultra-fast all-in-one FASTQ preprocessor. Bioinformatics 34: i884–i890

Cotticelli MG, Xia S, Truitt R, Doliba NM, Rozo AV, Tobias JW, Lee T, Chen J, Napierala JS, Napierala M, et al (2023) Acute frataxin knockdown in induced pluripotent stem cell-derived cardiomyocytes activates a type I interferon response. Dis Model Mech 16

Doncheva NT, Morris JH, Gorodkin J & Jensen LJ (2019) Cytoscape StringApp: Network Analysis and Visualization of Proteomics Data. J Proteome Res 18: 623–632

Edelstein AD, Tsuchida MA, Amodaj N, Pinkard H, Vale RD & Stuurman N (2014) Advanced methods of microscope control using μManager software. J Biol Methods 1

Figueira TR, Melo DR, Vercesi AE & Castilho RF (2012) Safranine as a fluorescent probe for the evaluation of mitochondrial membrane potential in isolated organelles and permeabilized cells. Methods Mol Biol 810: 103–117

Formisano L, Guida N, Valsecchi V, Pignataro G, Vinciguerra A, Pannaccione A, Secondo A, Boscia F, Molinaro P, Sisalli MJ, et al (2013) NCX1 is a new rest target gene: role in cerebral ischemia. Neurobiol Dis 50: 76–85

Frangogiannis N (2020) Transforming growth factor-β in tissue fibrosis. J Exp Med 217: e20190103

Fromigué O, Haÿ E, Barbara A & Marie PJ (2010) Essential role of nuclear factor of activated T cells (NFAT)-mediated Wnt signaling in osteoblast differentiation induced by strontium ranelate. J Biol Chem 285: 25251–25258

Gibb AA, Lazaropoulos MP & Elrod JW (2020) Myofibroblasts and Fibrosis: Mitochondrial and Metabolic Control of Cellular Differentiation. Circ Res 127: 427–447

Guida N, Serani A, Sanguigno L, Mascolo L, Cuomo O, Fioriniello S, Marano D, Ragione FD, Anzilotti S, Brancaccio P, et al (2024) Stroke Causes DNA Methylation at Heart Promoter in the Brain Via DNMT1/MeCP2/REST Epigenetic Complex. J Am Heart Assoc 13: e030460

Gu Z, Eils R & Schlesner M (2016) Complex heatmaps reveal patterns and correlations in multidimensional genomic data. Bioinformatics 32: 2847–2849

Hobai IA & O’Rourke B (2004) The potential of Na+/Ca2+ exchange blockers in the treatment of cardiac disease. Expert Opin Investig Drugs 13: 653–664

Hörtenhuber M, Toledo EM, Smedler E, Arenas E, Malmersjö S, Louhivuori L & Uhlén P (2017) Mapping genes for calcium signaling and their associated human genetic disorders. Bioinformatics 33: 2547–2554

Hudecova S, Lencesova L, Csaderova L, Sirova M, Cholujova D, Cagala M, Kopacek J, Dobrota D, Pastorekova S & Krizanova O (2011) Chemically mimicked hypoxia modulates gene expression and protein levels of the sodium calcium exchanger in HEK 293 cell line via HIF-1α. Gen Physiol Biophys 30: 196–206

Janssen LJ, Mukherjee S & Ask K (2015) Calcium Homeostasis and Ionic Mechanisms in Pulmonary Fibroblasts. Am J Respir Cell Mol Biol 53: 135–148

Koeppen AH, Becker AB, Feustel PJ, Gelman BB & Mazurkiewicz JE (2016a) The significance of intercalated discs in the pathogenesis of Friedreich cardiomyopathy. J Neurol Sci 367: 171–176

Koeppen AH, Qian J, Travis AM, Sossei AB, Feustel PJ & Mazurkiewicz JE (2020) Microvascular pathology in Friedreich cardiomyopathy. Histol Histopathol 35: 39–46

Koeppen AH, Ramirez RL, Becker AB & Mazurkiewicz JE (2016b) Dorsal root ganglia in Friedreich ataxia: satellite cell proliferation and inflammation. Acta Neuropathol Commun 4: 46

Kumari D & Usdin K (2009) Chromatin remodeling in the noncoding repeat expansion diseases. J Biol Chem 284: 7413–7417

Legeay M, Doncheva NT, Morris JH & Jensen LJ (2020) Visualize omics data on networks with Omics Visualizer, a Cytoscape App. F1000Res 9: 157

Li J, Rozwadowska N, Clark A, Fil D, Napierala JS & Napierala M (2019) Excision of the expanded GAA repeats corrects cardiomyopathy phenotypes of iPSC-derived Friedreich’s ataxia cardiomyocytes. Stem Cell Res 40: 101529

Li L-A, Wu Q-R, Luo L-B, Yang H, Liu H-Y, Li Q, Zhang M-Z, Deng C-Y, Rao F & Zhang Q-H (2026) Piezo1 Knockdown Attenuates Hypertension-Induced Cardiac Fibrosis by Inhibiting Ca/β-Catenin Signalling Pathway. Clin Exp Pharmacol Physiol 53: e70130

Lill R & Freibert S-A (2020) Mechanisms of Mitochondrial Iron-Sulfur Protein Biogenesis. Annu Rev Biochem 89: 471–499

Li Y, Lu Y, Polak U, Lin K, Shen J, Farmer J, Seyer L, Bhalla AD, Rozwadowska N, Lynch DR, et al (2015) Expanded GAA repeats impede transcription elongation through the FXN gene and induce transcriptional silencing that is restricted to the FXN locus. Hum Mol Genet 24: 6932–6943

Li Y, Polak U, Clark AD, Bhalla AD, Chen Y-Y, Li J, Farmer J, Seyer L, Lynch D, Butler JS, et al (2016) Establishment and Maintenance of Primary Fibroblast Repositories for Rare Diseases-Friedreich’s Ataxia Example. Biopreserv Biobank 14: 324–329

Lombardi AA, Gibb AA, Arif E, Kolmetzky DW, Tomar D, Luongo TS, Jadiya P, Murray EK, Lorkiewicz PK, Hajnóczky G, et al (2019) Mitochondrial calcium exchange links metabolism with the epigenome to control cellular differentiation. Nat Commun 10: 4509

Love MI, Huber W & Anders S (2014) Moderated estimation of fold change and dispersion for RNA-seq data with DESeq2. Genome Biol 15: 550

Lynch DR, Rojsajjakul T, Subramony SH, Perlman SL, Keita M, Mesaros C & Blair IA (2024) Frataxin analysis using triple quadrupole mass spectrometry: application to a large heterogeneous clinical cohort. J Neurol 271: 1844–1849

Manfredi G, Gupta N, Vazquez-Memije ME, Sadlock JE, Spinazzola A, De Vivo DC & Schon EA (1999) Oligomycin induces a decrease in the cellular content of a pathogenic mutation in the human mitochondrial ATPase 6 gene. J Biol Chem 274: 9386–9391

Marullo C, Croci L, Giupponi I, Rivoletti C, Zuffetti S, Bettegazzi B, Cremona O, Giunti P, Ambrosi A, Casoni F, et al (2025) Altered Ca2+ responses and antioxidant properties in Friedreich’s ataxia-like cerebellar astrocytes. J Cell Sci 138

McArthur L, Chilton L, Smith GL & Nicklin SA (2015) Electrical consequences of cardiac myocyte: fibroblast coupling. Biochem Soc Trans 43: 513–518

Miyara S, Adler M, Umansky KB, Häußler D, Bassat E, Divinsky Y, Elkahal J, Kain D, Lendengolts D, Ramirez Flores RO, et al (2025) Cold and hot fibrosis define clinically distinct cardiac pathologies. Cell Syst 16: 101198

Napierala JS, Li Y, Lu Y, Lin K, Hauser LA, Lynch DR & Napierala M (2017) Comprehensive analysis of gene expression patterns in Friedreich’s ataxia fibroblasts by RNA sequencing reveals altered levels of protein synthesis factors and solute carriers. Dis Model Mech 10: 1353–1369

Nurk S, Koren S, Rhie A, Rautiainen M, Bzikadze AV, Mikheenko A, Vollger MR, Altemose N, Uralsky L, Gershman A, et al (2022) The complete sequence of a human genome. Science 376: 44–53

Patro R, Duggal G, Love MI, Irizarry RA & Kingsford C (2017) Salmon provides fast and bias-aware quantification of transcript expression. Nat Methods 14: 417–419

Payne RM (2022) Cardiovascular Research in Friedreich Ataxia: Unmet Needs and Opportunities. JACC Basic Transl Sci 7: 1267–1283

Payne RM, O’Connell TM, Pride PM, Wagner GR, Eckert GJ, Johnson TR, Shou W & Hutchins GD (2025) Positron emission tomography reveals increased myocardial glucose uptake in a subset of Friedreich ataxia patients. Sci Rep 15: 37247

Piersma B, Wouters OY, de Rond S, Boersema M, Gjaltema RAF & Bank RA (2017) Ascorbic acid promotes a TGF1-induced myofibroblast phenotype switch. Physiol Rep 5

Posit team (2026) RStudio: Integrated Development Environment for R Posit Software, PBC

Raman SV, Phatak K, Hoyle JC, Pennell ML, McCarthy B, Tran T, Prior TW, Olesik JW, Lutton A, Rankin C, et al (2011) Impaired myocardial perfusion reserve and fibrosis in Friedreich ataxia: a mitochondrial cardiomyopathy with metabolic syndrome. Eur Heart J 32: 561–567

Reetz K, Lischewski SA, Dogan I, Didszun C, Pishnamaz M, Konrad K, Marx-Schütt K, Farmer J, Lynch DR, Corben LA, et al (2025) Friedreich’s ataxia-a rare multisystem disease. Lancet Neurol 24: 614–624

Reimand J, Isserlin R, Voisin V, Kucera M, Tannus-Lopes C, Rostamianfar A, Wadi L, Meyer M, Wong J, Xu C, et al (2019) Pathway enrichment analysis and visualization of omics data using g:Profiler, GSEA, Cytoscape and EnrichmentMap. Nat Protoc 14: 482–517

Romero JR, Rivera A, Lança V, Bicho MDP, Conlin PR & Ricupero DA (2005) Na+/Ca2+ exchanger activity modulates connective tissue growth factor mRNA expression in transforming growth factor beta1- and Des-Arg10-kallidin-stimulated myofibroblasts. J Biol Chem 280: 14378–14384

Rötig A, de Lonlay P, Chretien D, Foury F, Koenig M, Sidi D, Munnich A & Rustin P (1997) Aconitase and mitochondrial iron-sulphur protein deficiency in Friedreich ataxia. Nat Genet 17: 215–217

Saliba Y, Jebara V, Hajal J, Maroun R, Chacar S, Smayra V, Abramowitz J, Birnbaumer L & Farès N (2019) Transient Receptor Potential Canonical 3 and Nuclear Factor of Activated T Cells C3 Signaling Pathway Critically Regulates Myocardial Fibrosis. Antioxid Redox Signal 30: 1851–1879

Sayles NM, Napierala JS, Anrather J, Diedhiou N, Li J, Napierala M, Puccio H & Manfredi G (2023) Comparative multi-omic analyses of cardiac mitochondrial stress in three mouse models of frataxin deficiency. Dis Model Mech 16

Scognamiglio A, Corvino A, Caliendo G, Fiorino F, Perissutti E, Santagada V & Severino B (2025) Druggability of Sodium Calcium Exchanger (NCX): Challenges and Recent Development. Int J Mol Sci 26

Tai Y, Woods EL, Dally J, Kong D, Steadman R, Moseley R & Midgley AC (2021) Myofibroblasts: Function, Formation, and Scope of Molecular Therapies for Skin Fibrosis. Biomolecules 11

Tsou AY, Paulsen EK, Lagedrost SJ, Perlman SL, Mathews KD, Wilmot GR, Ravina B, Koeppen AH & Lynch DR (2011) Mortality in Friedreich ataxia. J Neurol Sci 307: 46– 49

Valsecchi V, Laudati G, Cuomo O, Sirabella R, Del Prete A, Annunziato L & Pignataro G (2021) The hypoxia sensitive metal transcription factor MTF-1 activates NCX1 brain promoter and participates in remote postconditioning neuroprotection in stroke. Cell Death Dis 12: 423

Vazana-Netzarim R, Elmalem Y, Sofer S, Bruck H, Danino N & Sarig U (2023) Distinct HAND2/HAND2-AS1 Expression Levels May Fine-Tune Mesenchymal and Epithelial Cell Plasticity of Human Mesenchymal Stem Cells. Int J Mol Sci 24

Wang J, Zhang Z, Hu Y, Hou X, Cui Q, Zang Y & Wang C (2007) SEA0400, a novel Na+/Ca2+ exchanger inhibitor, reduces calcium overload induced by ischemia and reperfusion in mouse ventricular myocytes. Physiol Res 56: 17–23

Wu P, Zhang Z, Zheng K, Zhu Z, Zhu Y, Kiram A, Zhao L, Chen H, Xu Z, Li X, et al (2026) RUNX2 Activation in Fibro/Adipogenic Progenitors Promotes Muscle Fibrosis in Muscular Dystrophy. Adv Sci (Weinh) 13: e10850

Wu T, Hu E, Xu S, Chen M, Guo P, Dai Z, Feng T, Zhou L, Tang W, Zhan L, et al (2021) clusterProfiler 4.0: A universal enrichment tool for interpreting omics data. Innovation (Camb) 2: 100141

Zhang B, Jiang J, Yue Z, Liu S, Ma Y, Yu N, Gao Y, Sun S, Chen S & Liu P (2016) Store-Operated Ca Entry (SOCE) contributes to angiotensin II-induced cardiac fibrosis in cardiac fibroblasts. J Pharmacol Sci 132: 171–180

Zhang H, Tian L, Shen M, Tu C, Wu H, Gu M, Paik DT & Wu JC (2019) Generation of Quiescent Cardiac Fibroblasts From Human Induced Pluripotent Stem Cells for In Vitro Modeling of Cardiac Fibrosis. Circ Res 125: 552–566

